# A metal-dependent switch moderates activity of the hexameric M17 aminopeptidases

**DOI:** 10.1101/244665

**Authors:** Nyssa Drinkwater, Wei Yang, Blake T. Riley, Brooke K. Hayes, Komagal Kannan Sivaraman, Tess R. Malcolm, Sarah C. Atkinson, Natalie A. Borg, Itamar Kass, Ashley M. Buckle, Sheena McGowan

## Abstract

The metal-dependent M17 aminopeptidases are conserved throughout all kingdoms of life. The large enzyme family is characterised by a conserved binuclear metal center and a distinctive homohexameric arrangement. To understand the mechanistic role of the hexameric assembly, we undertook an investigation of the structure and dynamics of the M17 aminopeptidase from *P. falciparum*, *Pf*A-M17. We describe a novel structure of *Pf*A-M17, which shows that the active sites of each trimer are linked by a dynamic loop, and that the loop movement is coupled with a drastic rearrangement of the binuclear metal center and substrate-binding pocket. Molecular dynamics simulations, supported by biochemical analyses of *Pf*A-M17 variants, demonstrate that this rearrangement is inherent to *Pf*A-M17, and that the transition between the active and inactive states is part of a dynamic regulatory mechanism. Key to the mechanism is a re-modelling of the binuclear metal center, which occurs in response to a signal from the neighbouring active site, and serves to moderate the rate of proteolysis under different environmental conditions. Therefore, this work has identified the precise mechanism by which oligomerization contributes to *Pf*A-M17 function. Further, it has described a novel role for metal cofactors in the regulation of enzymes with implications for the wide range of metalloenzymes that operate via a two-metal ion catalytic center including DNA processing enzymes and metalloproteases.

## Introduction

Intracellular proteolysis requires precise spatial and temporal control to prevent cleavage of proteins not destined for destruction. For this purpose, high-molecular weight protease enzymes are self-compartmentalised, whereby the proteolytic active sites are enclosed in inner cavities isolated from the cellular environment. Such an arrangement is often mediated by multimeric self-association, such as that seen for the family of M17 aminopeptidases, otherwise known as PepA or LAP (leucyl aminopeptidases; Clan MF, family M17) (MEROPS (1)), which are present in all kingdoms of life. Although overall sequence conservation is low, M17 aminopeptidases possess a conserved homohexameric structure wherein a dimer of trimers encloses an inner cavity harbouring the six active sites (2). The sequence and structure of the proteolytic sites themselves, as well as the reaction they catalyze, are highly conserved, all utilising two divalent metal ion cofactors to catalyze the removal of select N-terminal amino acids from short peptide chains (2). This proteolysis reaction contributes to intracellular protein turnover, a fundamental housekeeping process across all living organisms (3).

While M17 aminopeptidases are well known for their proteolytic roles, a wide range of additional functions beyond aminopeptidase activity have also been attributed to M17 family members (4). In bacteria, M17 aminopeptidases bind DNA in a sequence-specific manner to regulate transcription (5), and also form hetero-oligomers (6-8) which control site-specific DNA recombination (9, 10). In plants, M17 aminopeptidases function as part of the stress response (11) that, in addition to aminopeptidase activity (12), has been suggested to result from molecular chaperone activity (13). Furthermore, the molecular chaperone function has been attributed to a monomeric form of enzyme, suggesting that the hexamer can dissociate into individual protomers to moderate distinct functionalities (13, 14). Therefore, although the family of M17 aminopeptidases have a highly conserved structure across different organisms, they are multifunctional, capable of performing diverse organism-specific functions far beyond peptide hydrolysis (4). These diverse functions are largely mediated by dynamic changes in macromolecular assemblies; the conserved hexameric structure clearly has an enormous capacity to moderate functionality through structural dynamics. Such dynamics are also likely to influence the mechanism of proteolysis characteristic to the enzyme family. Preliminary analysis of the tomato M17 aminopeptidase (LAP-A1) has suggested that hexamerisation is required for proteolysis (15). However, crystal structures of M17 aminopeptidases from a wide range of organisms all show that each of the six active sites is entirely self-contained. The hexameric assembly of M17 aminopeptidases clearly plays an important role in proteolysis. Despite this, our mechanistic understanding of M17 proteolysis is based on early examinations of the bovine lens M17 aminopeptidase (blLAP), which treat the six chains within the hexamer as discrete entities, and have only ever considered small-scale flexibility within a single active site (16).

Of interest to our team is the M17 aminopeptidase from *Plasmodium falciparum* (*Pf*A-M17), the major causative agent of malaria in humans. *Pf*A-M17 is essential for the blood stage of the parasite life cycle (17) and inhibition of *Pf*A-M17 results in parasite death both *in vitro* (18) and *in vivo* (19). Consequently, *Pf*A-M17 is an exciting target for the development of novel antimalarials (19-24). The crystal structure of *Pf*A-M17 shows the homohexameric arrangement characteristic of M17 aminopeptidase enzymes (Fig. 1A), with the six active sites orientated inwards and accessible to the central cavity (Fig. 1B and C) (25). Herein, we examined the role hexamerisation plays in the *Pf*A-M17 mechanism through molecular dynamics (MD) simulations, X-ray crystallography, and mutational and solution-state analyses. With this comprehensive strategy, we characterised the range of protein motions inherent to *Pf*A-M17, and probed the specific contribution of those motions to enzyme function. Based on the results, we propose a novel model for how *Pf*A-M17 functions at an atomic level. In the interior of the catalytic cavity, six dynamic, regulatory loops (denoted L13), one contributed from each protomer, function cooperatively to mediate communication between the subunits of the hexamer. This loop motion shuttles a key conserved lysine residue, Lys386 between neighbouring active sites and promotes re-arrangement of the binuclear metal center. Based on mutagenesis and biochemical analyses, we propose a novel role for Lys386 in stabilization of the hexameric assembly through second-shell coordination interactions with the binuclear center, as well as shuttle of the substrate/product in/out of the active site. These roles for Lys386 are in addition to the previously proposed role of catalytic base in the reaction mechanism and stabilization of the substrate throughout the reaction. Further, we show that the L13 loop motion coupled with re-arrangement of the binuclear metal center effects a switch between active and inactive enzyme states, thereby acting as a dynamic regulatory mechanism. Based on the evidence, we propose this metal-dependent switch is an inherent component of the *Pf*A-M17 catalytic cycle, and is the key physiological mechanism that regulates cellular proteolysis under different environmental conditions.

**Figure 1.**
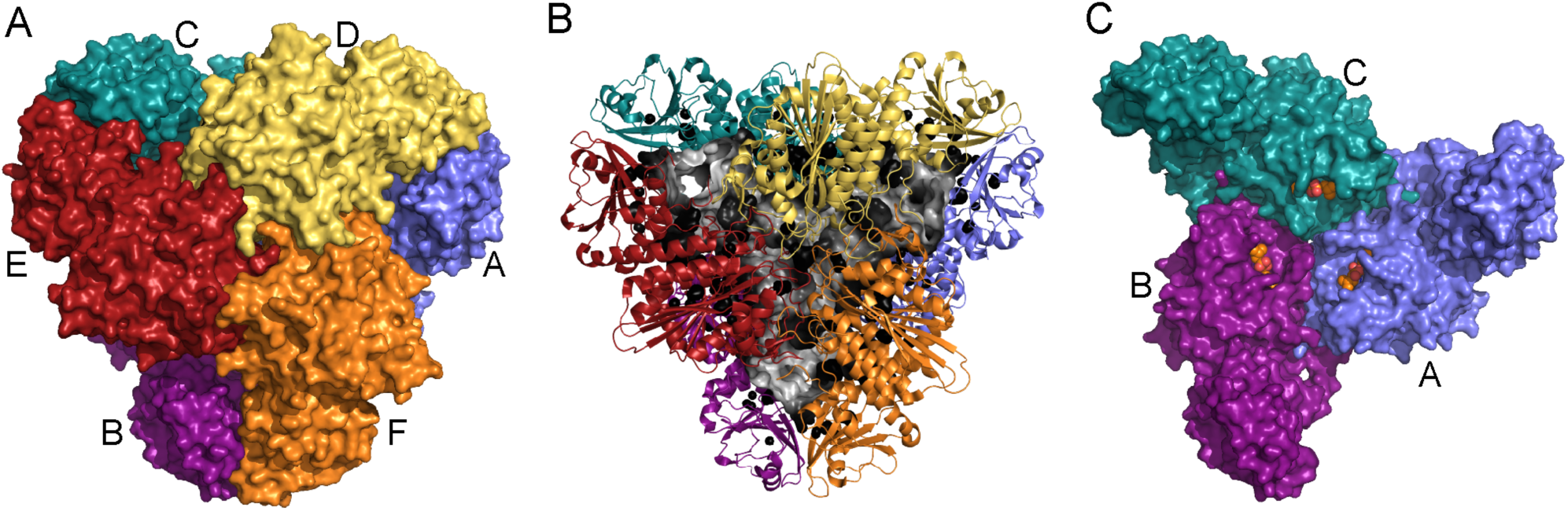
The crystal structure of *Pf*A-M17. (A) *Pf*A-M17 is a homo hexamer composed of a dimer of trimers. Three chains, denoted A (blue), B (purple), and C (teal) interact through their C-terminal domains to form a ‘trimer’, which interacts with an identical trimer made up of D (yellow), E (red), and F (orange) to form the hexamer. (B) *Pf*A-M17 (cartoon) possesses a large internal cavity (grey surface representation) that contains the six active sites. (C) A single trimeric face (ABC) of the hexamer, shown in the same orientation as A (front trimeric face hidden). Orange spheres show binding of substrate-analogue bestatin, and indicate the active sites, which are exposed to the inner cavity. (RCSB PDB ID for *Pf*A-M17, 3KQZ) (25).

## Materials and Methods

### Molecular biology, protein production and purification

The cloning of the truncated *Pf*A-M17 gene (encoding amino acids 85 – 605) into pTrcHis2B has been previously reported (25). Site directed mutagenesis was performed by PCR and confirmed by DNA sequencing, while deletion mutants were synthesised and sub-cloned into pET-21d(+) (GenScript). *Escherichia coli* strains DH5α and K-12 were used for DNA manipulation, and BL21(DE3) used for protein expression. Recombinant His_6_-tagged *wild type* and mutant *Pf*A-M17 were expressed using an autoinduction method as previously described for *wild type Pf*A-M17 (25). Proteins were expressed and purified as has previously been described for *wild type Pf*A-M17 (25) using a two-step purification procedure of Ni–NTA agarose column followed by gel filtration chromatography on a Superdex S200 10/300 column in 50 mM HEPES pH 8.0, 150 mM NaCl buffer.

### Protein analysis by analytical ultracentrifugation, circular dichroism spectroscopy, and gel filtration chromatography

Analytical ultracentrifugation (AUC) experiments were conducted using a Beckman Coulter Optima Analytical Ultracentrifuge at a temperature of 20 °C. For sedimentation velocity experiments, samples were loaded at a concentration of 0.5 mg/ml. 380 μL of sample and 400 μL of reference solution (50 mM HEPES pH 8.0, 1 mM MnCl_2_, 300 mM NaCl) were loaded into a conventional double sector quartz cell and mounted in a Beckman 4-hole An-60 Ti rotor. Samples were centrifuged at a rotor speed of 40,000 rpm and the data was collected continuously at 238 nm. Solvent density (1.0144 g/mL at 20 °C) and viscosity (1.0611 cP at 20 °C), as well as estimates of the partial specific volume (0.7419 mL/g for *Pf*A-M17 at 20 °C) were computed using the program SEDNTERP (26). Sedimentation velocity data were fitted to continuous size [*c*(*s*)] and continuous molar mass [*c*(*M*)] distribution models using the program SEDFIT (27, 28). For sedimentation equilibrium experiments, reference and sample sectors were loaded with 140 μL reference and 100 μL sample (0.5 mg/ml, 0.25 mg/ml and 0.1 mg/mL) plus 20 μL FC-43 oil in the sample sector. After initial scans at 3,000 rpm to determine optimum wavelength and radial range for the experiment, samples were centrifuged at 12,000, 18,000 and 40,000 rpm at 20 °C. Data were collected at multiple wavelengths every hour until sedimentation equilibrium was attained (24 h). Sedimentation equilibrium data were analysed with SEDPHAT (29) using the single species analysis model.

Circular dichroism (CD) measurements were performed using a Jasco 815 spectropolarimeter. Far-UV scans were performed at 190 – 250 nM, using 0.2 mg/mL protein concentration in PBS pH 7.5 and quartz curvette of 1 mm path length. Analytical gel filtration chromatography was performed on a Superdex S200 Increase 10/300 in 50 mM HEPES pH 8.0, 1 mM MnCl_2_, 150 mM NaCl. Approximate molecular weight of eluate was calculated by interpolation of a standard curve constructed with appropriate molecular weight standards.

### Aminopeptidase assays and analysis

Aminopeptidase activity was determined by measuring the release of the fluorogenic-leaving group, *L*-Leucine-7-amido-4-methylcoumarin hydrochloride (Sigma-Aldrich, L2145) (H-Leu-NH-Mec), from the fluorogenic peptide substrates H-Leu-NHMec as described previously (30). Briefly, assays were performed in triplicate and carried out in 50 μL total volume in 100 mM Tris-HCl, pH 8.0, 1 mM MnCl_2_ at 37 °C and activity was monitored until steady-state was achieved. Fluorescence was measured using a FluoroStar Optima plate reader (BMG Labtech), with excitation and emission wavelengths of 355 nm and 460 nm, respectively. Activity of the association mutant *Pf*A-M17(W525A, Y533A) was assessed with 10 μM concentration of substrate H-Leu-NH-Mec and compared to *wild type Pf*A-M17 activity assessed under identical reaction conditions.

For active enzymes, the Michaelis constant, *K*_M_, was calculated from the initial rates over a range of substrate concentrations (H-Leu-NH-Mec, 0.5 – 500 μM) with enzyme concentrations fixed at 40 nM for *wild type Pf*A-M17, 10 nM for *Pf*A-M17(D394A), 1000 nM for *Pf*A-M17(K386A), 650 nM for *Pf*A-M17(A387P) and *PfA*-M17(Δ388-389), and *Pf*A-M17(Δ388-390). Gain was fixed at 800 for all Michaelis-Menten assays, and converted to units of product by interpolation of a standard curve constructed (fluorescence output of a range of methylcoumarin concentrations, collected at gain of 800). Kinetic parameters including, *K_M_*, *k_cat_*, and *n*_H_ were calculated with non-linear regression protocols by using GraphPad Prism 7.

### Crystallization, data collection, structure determination and refinement

*Pf*A-M17 was concentrated to 10 mg/mL in 50 mM HEPES pH 8.0, 150 mM NaCl for crystallization. Crystals were grown by hanging drop vapour diffusion in 20 % PEG3350, 0.2 M calcium acetate, with drops composed of 2 μL protein plus 1 μL precipitant. Crystals grew to large plates in 7 days. Crystals were cryoprotected in mother liquor supplemented with 15 % 2-methyl-2,4-pentanediol for 30 s before flash cooling in liquid nitrogen.

Data were collected at 100 K using synchrotron radiation at the Australian Synchrotron using the micro crystallography MX2 beamline 3ID1 (31). Data were processed using XDS (32) and scaled in Aimless (33) as part of the ccp4 suite (34). The structure was solved by molecular replacement in Phaser using the structure of unliganded *Pf*A-M17 (RCSB ID 3KQZ) as the search model (25). Refinement was carried out with iterative rounds of model building in Coot (35, 36) and refinement using Phenix (37). Water molecules and metal ions were placed manually based on the presence of *F*_o_–*F*_c_ and 2*F*_o_–*F*_c_ electron density of appropriate signal. To determine metal ion identity, X-ray diffraction data was collected above and below the Zn^2+^ anomalous absorption edge, and used to calculate anomalous maps. If anomalous density was not visible below a 3.0 σ contour level, the metal was modelled as an Mn^2+^, as determined by intensity level of the *F*_o_–*F*_c_ and 2*F*_o_–*F*_c_ electron density maps.

The structure was validated with MolProbity (38) and figures generated using PyMOL version 1.8.23. Final structure coordinates were deposited in the protein databank (RCSB ID 6NWI), and a summary of data collection and refinement statistics is provided in Supplementary Information (Supp. Info. 7). For clarity and consistency, the chains in the deposited structure are numbered according to the model of unliganded *Pf*A-M17 (RCSB PDB ID 3KQZ).

### MD system setup and simulation protocol

The starting *Pf*A-M17 model for MD simulations was based on the unliganded crystal structure (RCSB PDB ID 3KQZ. To prepare the model, missing atoms and residues (A^84, 257-261^, B^84-85, 255-262^, C^84-85, 255-259^, D^84, 255-259^, E^84-85, 152^, F^84-85, 136, 255-261^) were rebuilt using Modeller v9.11 (39), protonated according to their states at pH 7.0 using the PDB2PQR server (40) and subjected to energy minimization using Modeller (39). The hexameric *Pf*A-M17 system consisted of ∼ 294,000 atoms with a periodic box of 167 Å × 166 Å × 121 Å and the monomeric *Pf*A-M17 system consisted of ∼ 81,000 atoms with a periodic box of 120 Å × 85 Å × 94 Å (after solvation). Periodic boundary conditions (pbc) were used for all simulations. System charges were neutralised with sodium counter ions. Proteins and ions were modelled using the AMBER force field FF12SB (41), the metal center was defined as described previously (42) and waters represented using the 3-particle TIP3P model (43). All atom MD simulations were performed using NAMD 2.9 on an IBM Blue Gene/Q cluster (monomer simulations) or x86 (hexamer simulations). Equilibration was performed in three stages. First, potential steric clashes in the initial configuration were relieved with 50,000 steps of energy minimisation. Initial velocities for each system were then assigned randomly according to a Maxwell–Boltzmann distribution at 100 K. Each system was then heated to 300 K over 0.1 ns, under the isothermal-isometric ensemble (NVT) conditions, with the protein atoms (excluding hydrogens) harmonically restrained (with a force constant of 10 kcal mol^−1^ A^−2^). Following this, each system was simulated for 100 ps under the isothermal-isobaric ensemble (NPT) with heavy atoms restrained. The harmonic restraints used were reduced from 10 to 2 kcal mol^−1^ A^−2^ during the simulations. The above equilibration process was performed three times from the same starting structure in order to initiate three production simulations with different initial velocities. For production simulations, the time step was set to 2 fs and the SHAKE algorithm was used to constrain all bonds involving hydrogen atoms. All simulations were run at constant temperature (300 K) and pressure (1 atm), using a Langevin damping coefficient of 0.5 fs^−1^, and a Berendsen thermostat relaxation time of τ_P_ = 0.1 ps. The Particle-Mesh Ewald (PME) method was used to set the periodic boundary conditions (PBC) that were used for long-range electrostatic interactions and a real space cut-off of 10 Å was used. Conformations were sampled every 10 ps for subsequent analysis. All frames with time interval of 10 ps were saved to disk.

### MD Analysis

Simulation trajectories were analysed using the GROMACS 5.14 simulation package. For principle component analysis (PCA), 3N*3N atom covariance matrices of the protein displacement in simulations were generated based on backbone atoms (N, Cα, C, O) of the *Pf*A-M17 crystal structure. Principle components (PCs) that taken together accounted for more than 50 % of the overall covariance, were chosen for essential dynamics analysis. The GROMACS 5.14 simulation package was used to project the trajectory onto the top PCs. Graphs and plots were produced with Xmgrace and GraphPad Prism7. Molecular graphics were prepared with PyMOL 1.8.23 and VMD1.9.3.

## Results

### Hexamerization is essential for catalytic function

Previously determined crystal structures of M17 aminopeptidases, including *Pf*A-M17 (Fig. 1) show that the six active sites are structurally distinct (2, 25). Each of the active sites is composed of residues from a single chain and contains its own catalytic machinery that includes a binuclear metal center and a carbonate ion (2). Based on current evidence, the hexameric assembly has no apparent role in enzyme function. To probe the role of hexamerisation in catalysis, we first attempted to identify whether any cooperativity between of substrate binding exists, which could indicate communication between the active sites. We performed substrate saturation experiments, wherein we assessed the relationship between substrate concentration and reaction velocity to analyse the Hill’s coefficient (*n*_H_). Using a fluorescently labelled substrate in the presence of 1 mM metal cofactor (manganese, Mn^2+^), we were unable to detect any evidence of cooperativity in *Pf*A-M17 catalysis (*n*_H_ = 1.1 ± 0.1, Fig. 2A). In the absence of evidence suggesting communication between the active sites, we questioned whether the *Pf*A-M17 hexamer was genuinely required for aminopeptidase activity. We addressed this question directly through the design of amino acid mutations to disrupt hexamerization.

**Figure 2.**
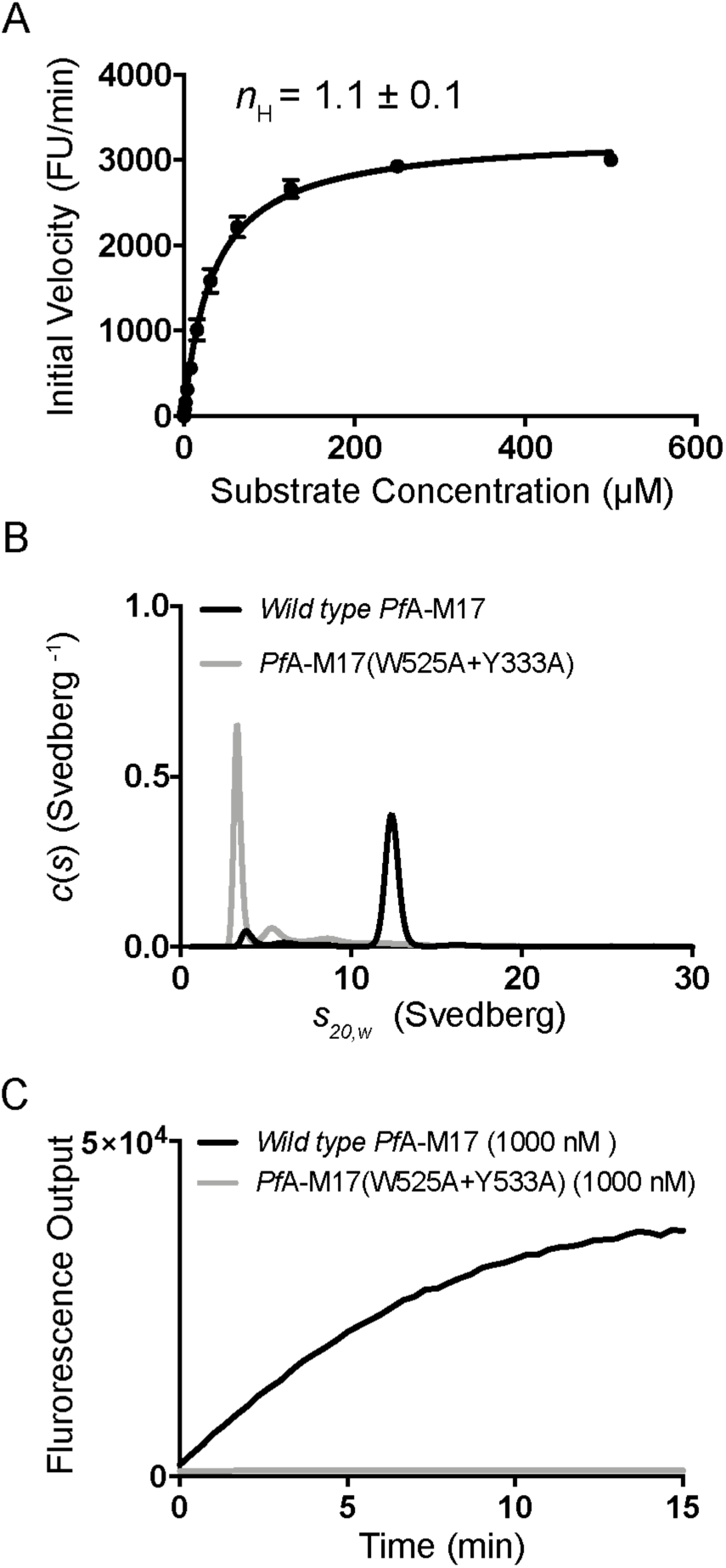
Hexameric arrangement of *Pf*A-M17 required for proteolysis. (A) Analysis of the Hills coefficient (*n*_H_) by substrate-saturation experiment showed no evidence of cooperativity in *Pf*A-M17 catalysis. Data shown are means ± SD (*n* = 3). (B) Sedimentation velocity profiles of *wild type Pf*A-M17 (black) and *Pf*A-M17(W525A+Y533A) (grey) in 1.0 mM MnCl_2_ and represented as continuous *c*(*s*) distribution (complete data shown in Supplementary Figure 2). (C) Assessment of the catalytic activity of *Pf*A-M17(W525A+Y533A) (grey) under *wild type* (black) conditions, showed that the monomeric enzyme is inactive.

Examination of the *Pf*A-M17 crystal structure identified Trp525 and Tyr533 as key residues that mediate association of each of the ‘trimers’ into the hexamer. From this, we hypothesised that their elimination should result in the formation of a stable trimer. The mutant *Pf*A-M17(W525A+Y533A) expressed and purified similarly to *wild type* enzyme, and analysis by circular dichroism (CD) showed that both mutant and *wild type* enzymes possess similar secondary structure elements (Supp. Info. 1). To investigate the oligomeric state of *Pf*A-M17(W525A+Y533A), we performed sedimentation velocity experiments. *c*(*s*) analysis showed the mutated enzyme sedimented as a predominantly single species with sedimentation coefficient 3.3S. In contrast, *wild type Pf*A-M17 was a majority 12.4S species, with a smaller, secondary species also observed (Fig. 2B, Supp. Info. 2A and 2B). Subsequent *c*(*M*) analysis suggested *Pf*A-M17(W525A+Y533A) was monomeric (Supp. Info. 2B), which was confirmed by sedimentation equilibrium experiments that determined a molecular weight of 59.4 kDa for *Pf*A-M17(W525A+Y533A) (Supp. Fig. 2C–E). We examined the activity of *Pf*A-M17(W525A+Y533A) (Fig. 1C), and determined that under *wild type* assay conditions, the monomeric enzyme was inactive, even at high concentrations (Fig. 1C). Therefore, hexamer formation is clearly essential for *Pf*A-M17 proteolytic activity and it appears that an M17 trimer is not stable in solution.

### Hexameric assembly stabilizes and links the *Pf*A-M17 active sites

Having established that monomeric *Pf*A-M7 is unable to function independently, we continued to search for the role of hexamerization. We turned to all-atom molecular dynamics (MD) simulations to probe the motions inherent to the *Pf*A-M17 hexamer, and additionally, compared the motions of the active hexamer to the inactive monomer. We performed all-atom MD simulations of both arrangements in triplicate (hexamer, 3 × 400 ns; monomer, 3 × 200 ns). Analysis of the root mean square deviation (RMSD) of the C*α* atoms over the course of the simulations indicated that the monomer underwent greater structural changes than the hexameric enzyme (average RMSD of monomer = 4.1 ± 0.5 Å, Supp. Info. 3A; average RMSD of hexamer = 2.5 ± 0.04 Å, Supp. Info. 3B). Further, in the simulation of monomeric *Pf*A-M17, variation was observed between triplicate runs (Supp. Info. 3A). This is in contrast to the hexamer simulation where all three runs showed a similar RMSD profile (Supp. Info. 3B). This clear difference in the dynamics of hexameric versus monomeric *Pf*A-M17 suggests that the hexameric assembly stabilizes the monomers to enable catalysis.

To determine how the stability of the hexamer translates to the catalytic machinery, we examined the local environment of the active sites throughout the simulation. The catalytic reaction mechanism of M17 aminopeptidases relies on the deprotonation of a catalytic water or hydroxyl ion for nucleophilic attack on the peptide carbonyl carbon (16, 44). In the simulation of monomeric *Pf*A-M17, we did not observe a stable position for a water molecule within the active site (Supp. Info. 4A). In contrast, in the simulation of hexameric *Pf*A-M17, we observed a stable water conformation wherein it associates with the site 1 Zn^2+^ ion (Supp. Info. 4B), consistent with the position of a nucleophilic water (16, 42).

Catalysis relies on precise placement of chemical moieties within the active site, so we assessed the mobility of individual residues throughout the simulations via calculation of the root mean square fluctuations (RMSF) of the Cα atoms. The RMSF analysis showed clear differences between the two simulations, particularly residues 385–391, which lie on a loop (referred to here as L13) flanking the active site on the interior of the central cavity. In the simulation of monomeric *Pf*A-M17, L13 exhibited a high degree of flexibility (average RMSF over three runs for L13 = 4.5 ± 0.4 Å; Supp. Info. 5A). However, in the simulation of hexameric *Pf*A-M17, the flexibility of the loop varied between the six chains (Supp. Info. 5B). In four chains (chain A, D, E, and F), L13 was relatively stable (average RMSF = 1.6 – 2.1 Å), however in the remaining two chains (chain B and C), it showed a higher level of mobility (average RMSF B = 2.8 ± 0.4 Å and C = 3.1 ± 0.4 Å).

To extract the details of the motions that the hexameric *Pf*A-M17 underwent, we performed a principal component analysis (PCA) of these simulations. Projecting the trajectories on to the top two PCs (where PC1 accounts for 62 % of the total variance and PC2 10 %) showed that the major motion, described by PC1, is an overall expansion of the hexamer (measured across the face of the trimer) from 120 Å to 127 Å (average of three measurements between Cα of Asn181*^a^*, Asn181*^b^* and Asn181*^c^*). The expansion is approximately 6 % of the overall hexamer size, and likely results primarily from the release of crystal constraints (45). However, within PC2 we observed a substantial movement of the L13 loop, the same region identified in the RMSF analysis. In the crystal structure, and therefore the starting conformation of the MD simulation, the position of L13 is identical in all six units, where it lines the active site and extends into the solvent of the inner cavity. PCA showed that in PC2 of the *Pf*A-M17 hexamer simulation, L13 from one subunit of each trimer (B and E) moves away from its own active site, extends across the entrance of the pocket and stretches toward the active site of the neighbouring chain (Supp. Info. 6). In the most extreme conformation in chain E, L13 occludes the entrance to the chain E pocket, and contacts L13 of the neighbouring chain, F. Although L13 shows some movement in all chains within the hexamer, the extreme motion is only observed for one site per trimer (chains B and E). Interestingly, L13 contains lysine residue 386, which has a predicted role in the reaction mechanism of *Pf*A-M17 (16, 44, 46). The loop movement therefore serves not only to link neighbouring active sites, but also to re-model the catalytic machinery. Taken together, our MD analyses demonstrates that the multimeric assembly stabilizes the active site environment of M17 aminopeptidases, and further, suggests that L13 might mediate communication between the active sites of the *Pf*A-M17 hexamer.

### Novel PfA-M17 conformation captured by crystallography

Our MD simulations identified that the L13 loop is dynamic and capable of linking one *Pf*A-M17 active site to the neighbouring unit within the trimer. This is the first evidence that shows the active sites of M17 aminopeptidases might be linked. However, MD simulations only provide a snapshot of the range of motion that proteins may experience, and indeed, due to the challenges of simulating metalloenzymes, may bias trajectories close to the active site (42). Therefore, we sought to validate our computational findings and demonstrate that the enzyme is physiologically capable of sampling different active site conformations.

We rationalised that meta-stable conformations of *Pf*A-M17, including variations of the position of L13, could be captured and characterised by crystallization. Therefore, we performed sparse matrix screens to identify novel *Pf*A-M17 crystallization conditions. Any conditions different to those from which the original crystal structure was determined were optimized, diffraction data collected, and structures solved by molecular replacement. One structure, solved to 2.0 Å (Supp. Info. 7), showed a previously unobserved conformation. Overall, the quaternary structure is similar to previous *Pf*A-M17 structures with the hexamer arranged as a dimer of trimers, and the domain arrangement in each unit consistent with previous conformations. However, the new structure shows a vastly different active site arrangement to any previously published M17 structure. In the previously determined *Pf*A-M17 structure, henceforth referred to as the ‘active’ conformation, L13 lines the active site and extends into the solvent of the inner cavity (Fig. 3A, 3B). In the new conformation, L13 loops of each of the six chains cross the entrance to the active sites and extend to the active site of the neighbouring chain in the trimer (Fig. 3C, 3D). This rearrangement is remarkably similar to those observed for chains B and E of the MD simulations. In this new position, the six key Lys386 residues each occupy the active site of the neighbouring chain (Fig. 3D). The L13 rearrangement therefore results in a direct link between the three active sites of each of the trimers within hexameric *Pf*A-M17. A relatively large movement of approximately 13 Å is required to elicit the conformational changes in each of the six loops, which also serves to occlude the entrances to the six sites (Fig. 4). To obtain the slack within L13 to adopt this extended, occluded conformation, the secondary structure at both ends of the loop has been disrupted (Fig. 3A compared to Fig. 3C, Fig. 4C). This includes disruption of the α-helical structure of ten residues (392–401), which results in substantial shortening of an active site helix (392–417), and complete disruption of a short *β*-strand (residues 372–379).

**Figure 3.**
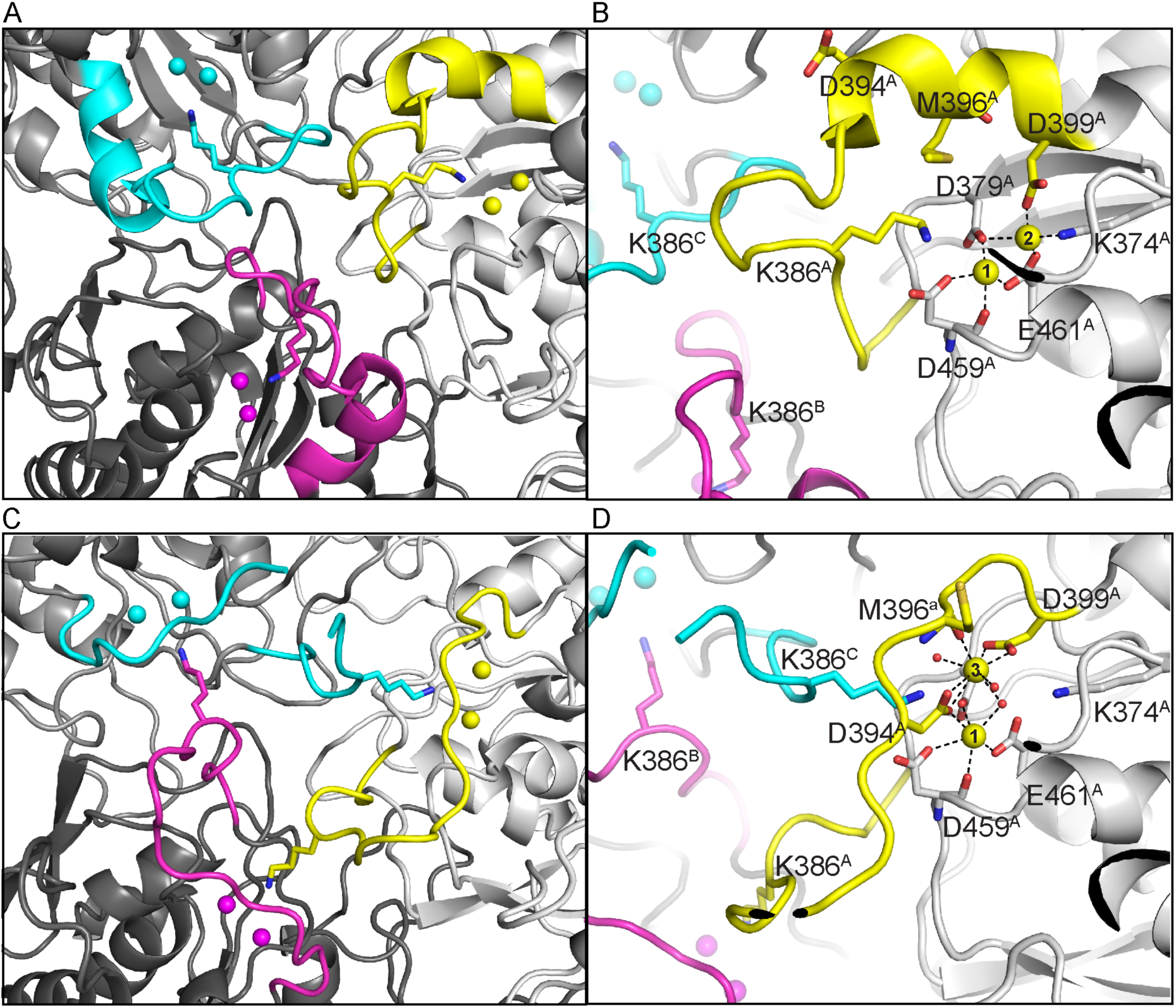
Novel *Pf*A-M17 structure captured with the active sites linked by flexible L13 loop. (A) Active site arrangement of active *Pf*A-M17 at the junction of the ABC trimer (on the interior of the *Pf*A-M17 hexamer) showing that the active sites of each chain are distinct. Chain A is light grey, with L13 loop and metal ions atoms in yellow; chain B is dark grey, with L13 loop and metal ions in magenta, chain C is medium grey, with L13 loop and metal ions in cyan. Lys386 of all chains shown in stick representation. (B) The active site of chain A in active *Pf*A-M17. Orientation is similar to that in panel A. Metal binding positions 1 and 2 are indicated, and active site residues shown in stick representation. Subscript denotes chain identifier. (C) Active site arrangement of inactive *Pf*A-M17 at the junction of the ABC trimer, showing that the active sites are linked by the L13 loops. Coloring and orientation consistent with panel A. Lys386 sits in the active site of the neighbouring chain, effectively linking all three active sites within the trimer. (D) Active site of chain A in inactive *Pf*A-M17 in same orientation as panel B. Disruption of active site α-helix is observed, and re-organisation of the active site architecture, including metal binding positions has occurred. Metal binding positions 1 and 3 are indicated, and active site residues shown in stick representation. (RCSB PDB ID, 6NWI).

**Figure 4.**
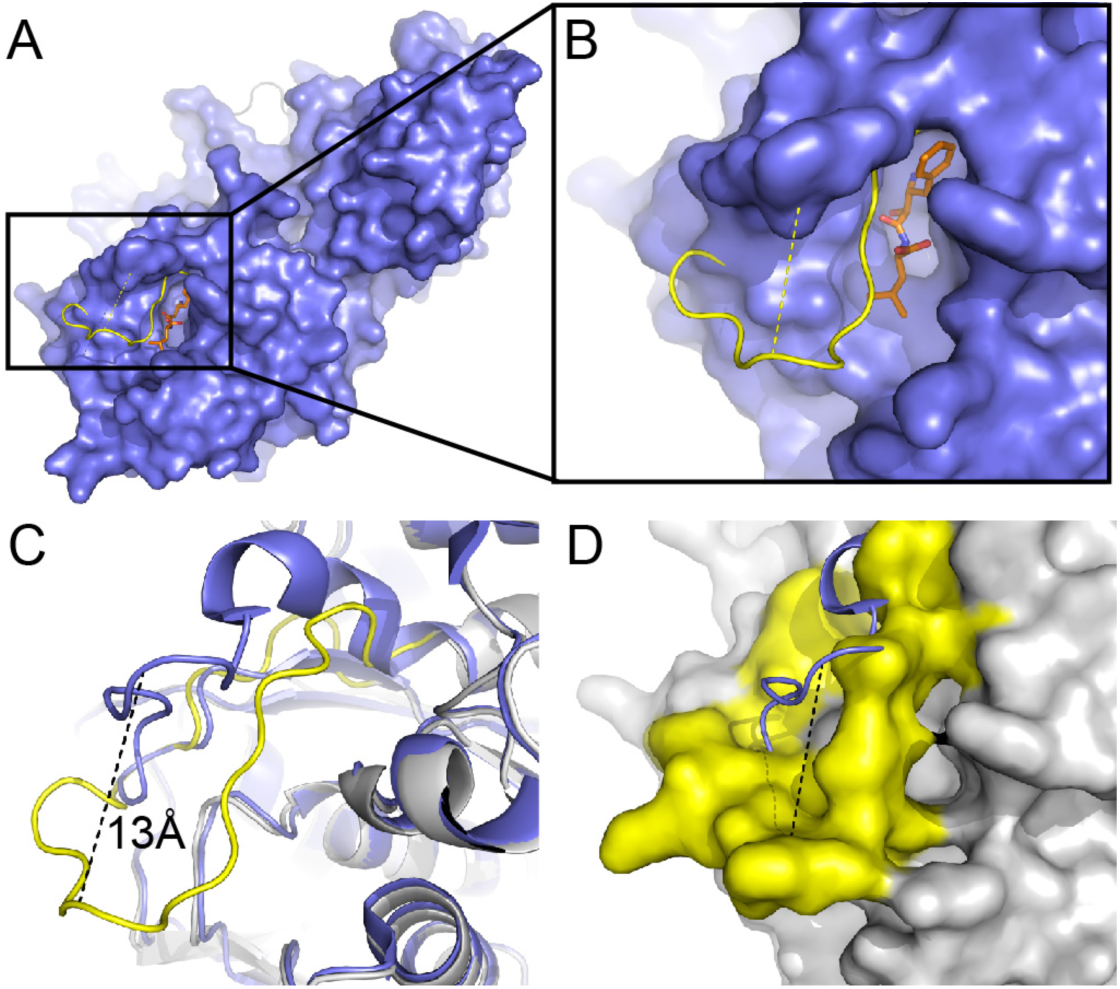
Conformational change from active to inactive *Pf*A-M17 mediated by flexible L13 loop. (A and B) Overlay of a single representative monomer (chain A) of the ‘inactive’ *Pf*A-M17 conformation (yellow cartoon) with ‘active’ *Pf*A-M17 (blue surface) in complex with substrate-analog bestatin (orange sticks). In the ‘active’ conformation (blue), the binding mode of bestatin (orange) is shown to indicate the active site. Yellow cartoon protruding through the surface shows the new position of the L13 loop in the ‘inactive’ conformation of *Pf*A-M17. The loop moves ∼13 Å (yellow dash) to occlude the active site, occupying the same space as bestatin in the ‘active’ structure. (C) Overlay of ‘active’ (blue cartoon) and ‘inactive’ (grey and yellow cartoon) crystal structures of *Pf*A-M17. Region and orientation is the same as that shown in B. Black dashed line indicates the ∼13 Å movement of the flexible L13 loop (yellow), which disrupts an active site alpha helix (top right, foreground) and beta sheet (top right, background). (D) Inverted representation of panel C, wherein the ‘inactive’ conformation of *Pf*A-M17 (grey) is shown in surface representation, with the flexible loop (yellow) completely occluding the active site. Black dash shows the 13 Å movement of the loop from the ‘active’ conformation (blue cartoon).

The active site rearrangement extends beyond protein backbone changes and includes the two catalytic metal ions (Supp. Info. 8 shows a morph between the two structures). Based on kinetic and biophysical characterisation, the two metal sites of the M17 aminopeptidases have previously been termed site 1 and site 2, whereby site 1 is that closest to the mouth of the active site (Fig. 3B) (2). Crystallographic studies show that site 1 metal ions are readily lost, while site 2 is always occupied (most often by Zn^2+^ or Mn^2+^ (4)), and kinetic studies have suggested that removal of the site 2 metal ion results in ablation of catalytic activity (47). Therefore, site 2 is generally considered the tight-binding, catalytic site, and site 1 the weak-binding, regulatory site (25, 47-49). Remarkably, in the novel conformation described here, we observed re-arrangement of the metal binding positions. While site 1 is occupied, the ‘catalytic’ metal in site 2 is absent (Fig. 3D). Further, electron density indicative of a metal ion is observed in a third, previously uncharacterised site ∼ 3.8 Å from site 1 and coordinated by the side chains of Asp394 and Asp399, the main chain oxygen of Met362, and two ordered water molecules (Fig 3D). Based on the octahedral coordination and a poor anomalous signal intensity at the Zn^2+^ edge, we modelled an Mn^2+^ ion into the density and termed the position site 3. In contrast, the metal bound to site 1 (capped trigonal bipyramidal coordination geometry) has a weak anomalous signal at the Zn^2+^ edge, but electron density could not always accommodate a fully occupied Zn^2+^ ion; we therefore suggest that site 1 contains a mix of bound Zn^2+^ and Mn^2+^, and have modelled Zn^2+^ at partial occupancy (0.8 – 1.0) where necessary. Site 3 is only available when L13 has undergone the described conformational change; the structure therefore shows a completely remodelled active site, including the architecture of the binding pocket and the catalytic binuclear center. This novel active site conformation is incompatible with binding substrate, so as a result, was termed the ‘inactive’ conformation.

The presence of a third metal binding site in the inactive conformation of *Pf*A-M17 was surprising since, to our knowledge, an equivalent site has never been observed in any M17 aminopeptidase. We were therefore curious to examine potential functional roles for the third metal ion. We identified Asp394 as crucial to the formation of site 3. This residue coordinates the site 3 metal ion and additionally, lies within the flexible L13 that has undergone substantial rearrangement (Fig. 3D). In contrast, in the original, active conformation of *Pf*A-M17, Asp394 has no clear role; while close to the active site, the side chain is directed to solvent and makes no interactions (Fig. 3B). Therefore, to validate a physiological role for the site 3 metal ion, we disrupted its ability to coordinate metal by mutating Asp394 to alanine, *Pf*A-M17(D394A). The effect of this mutation on enzyme activity was profound and unexpected. In comparison to *wild type* enzyme, *Pf*A-M17(D394A) showed greatly increased catalytic ability. The overall catalytic efficiency increased almost 10-fold, which resulted from an altered catalytic turnover rate (increased *k_cat_*, Table 1) rather than substrate binding affinity (*K_M_* largely unchanged, Table 1). Given Asp394 does not have a key structural role in the ‘active’ conformation of *Pf*A-M17, nor the currently accepted mechanism of hydrolysis, the change in activity observed on mutation of Asp394 could only result if the ‘inactive’ conformation has a functional role in *Pf*A-M17 catalysis, and is sampled as part of either the catalytic or regulatory mechanism.

**Table 1.** Kinetic analysis of mutant *Pf*A-M17 enzymes.

| | $[E]_{\text{tot}}$ ($\mu\text{M}$) | $K_M \pm \text{S.E.M}$<br>($\mu\text{M}$) | $k_{\text{cat}} \times 10^{-6} \pm \text{S.E.M.}$<br>($\text{min}^{-1}$) | $k_{\text{cat}} / K_M$ (M) |
| --- | --- | --- | --- | --- |
| *Wild type PfA*-M17 | 0.040 | $33.7 \pm 1.5$ | $2.5 \pm 0.03$ | 0.08 |
| *PfA*-M17(D394A) | 0.010 | $19.5 \pm 1.1$ | $11 \pm 0.2$ | 0.6 |
| *PfA*-M17(K386A) | 1.0 | $68.0 \pm 7.0$ | $0.052 \pm 0.002$ | 0.0008 |
| *PfA*-M17(A387P) | 0.65 | $45.1 \pm 2.5$ | $0.22 \pm 0.004$ | 0.005 |
| *PfA*-M17(Δ388–389) | 0.65 | $51.2 \pm 5.1$ | $0.31 \pm 0.01$ | 0.006 |
| *PfA*-M17(Δ388–390) | 0.080 | $102 \pm 9$ | $1.3 \pm 0.04$ | 0.01 |

### Dynamic loop regulates *Pf*A-M17 catalysis

The description of an inactive conformation of *Pf*A-M17, with active sites linked through the dynamic re-arrangement of L13, has clear mechanistic implications. Lys386, within the L13 loop, has a predicted role in the *Pf*A-M17 catalytic mechanism, where it is suggested to deprotonate the catalytic water and position the peptide substrate for catalysis. However, it is also a key player in the re-arrangement of L13 (Fig. 3B versus 3D). In the inactive conformation, Lys386 of each chain occupies the active site of the neighbouring chain. To probe the importance of Lys386 to the function of *Pf*A-M17, we mutated it to an alanine. *Pf*A-M17(K386A) showed substantially reduced activity compared to the *wild type* enzyme (Table 1). Determination of catalytic parameters showed this retardation of activity is largely due a decrease in the ability of the enzyme to accelerate the reaction (measured by a decrease in *k_cat_*, Table 1), but also to a slight reduction in the binding affinity of substrate (*K*M is 2-fold reduced compared to *wild type*, Table 1). While the disrupted activity demonstrates that Lys386 is key to *Pf*A-M17 function, it does not discriminate between a direct role in the catalytic reaction versus structural roles. To investigate further, we examined the affect of the K386A mutation on the solution state of *Pf*A-M17 using analytical gel filtration. The *wild type* enzyme elutes as a large molecular weight species with predicted molecular weight of 330 kDa (Supp. Info. 9A), consistent with the calculated molecular weight of a hexamer (350 kDa). In contrast, *Pf*A-M17(K386A) elutes as a smaller species with predicted molecular weight of 145 kDa (Supp. Info. 9B). Lys386 therefore plays a key role beyond catalysis that influences the stability of the hexameric assembly. To determine if Lys386 alone stabilizes the hexamer, or alternatively, if it is the motion of L13 shuttling Lys386 between neighbouring protomers, we created a series of L13 variants.

We first reduced the mobility of L13 by introducing a proline in place of Ala387, a loop residue that does not have any binding interactions in either the active or inactive conformations of *Pf*A-M17. Similarly to *Pf*A-M17(K386A), *Pf*A-M17(A387P) showed substantially reduced catalytic activity (Table 1). Large concentrations of enzyme were required to measure enzyme activity, which manifested as reduced product turnover (decreased *k_cat_*) compared to the *wild type* enzyme. However, unlike *Pf*A-M17(K386A), the hexameric assembly of *Pf*A-M17(A387P) is intact, as measured by analytical size exclusion chromatography (Supp. Info. 9C). Therefore, the ablation of activity is a direct result of the decrease in loop flexibility, not an alteration in oligomerization.

Finally, we shortened the loop by deletion of either two or three residues in *Pf*A-M17(Δ388–389) and *Pf*A-M17(Δ388–390). With these variants, we aimed to conserve the active site structure of the ‘active’ conformation, while preventing the linkage of neighbouring sites as observed in the ‘inactive’ conformation. Analytical size exclusion chromatography showed that neither deletion mutation compromised the stability of the hexamer (Supp. Info. 9D), yet both enzymes showed disrupted function. Deletion of only two residues in *Pf*A-M17(Δ388–389) reduced the catalytic turnover similarly to the *Pf*A-M17(A387P) mutation. However, deletion of an additional residue in *Pf*A-M17(Δ388–390) had a markedly different result. During assay optimization, we noted that *Pf*A-M17(Δ388–390) displayed highly variable and unpredictable levels of activity. With careful optimization, we were able to achieve reproducible kinetic parameters (Table 1); however, we chose to investigate the unusual activity profile further.

We compared the behaviour of *wild type Pf*A-M17 and *Pf*A-M17(Δ388–390) under different environmental conditions by simultaneously varying the concentrations of enzyme and Mn^2+^. At low concentrations of protein (125 – 250 nM), *wild type Pf*A-M17 demonstrated positive cooperativity with a Mn^2+^ cofactor, as measured by calculation of the Hill’s coefficient (*n*_H_ = 1.6 – 2.0, Table 2 and Supp. Info. 10A). Therefore, while our earlier results demonstrated that *Pf*A-M17 does not demonstrate cooperativity with the substrate, it does with metal cofactor. However, with increasing concentrations of enzyme (500 – 1000 nM), the cooperativity with the Mn^2+^ cofactor is lost (*n*_H_ = 0.71 – 1.2) (Table 2 and Supp. Info. 10A). Therefore, the mechanism by which *wild type Pf*A-M17 functions under different Mn^2+^ conditions is concentration dependent. In contrast, *Pf*A-M17(Δ388–390) only demonstrates positive cooperativity with the Mn^2+^ cofactor when the enzyme is at high concentrations ([E] = 1000 nM, *n*_H_ = 2.2, Table 2 and Supp. Info. 10B), while at low concentrations (≤ 500 nM), the enzyme shows negative cooperativity (*n*_H_ = 0.53 – 0.63, Table 2 and Supp. Info. 10B). Together, this data demonstrates that L13, while not absolutely required for enzyme function, mediates the mechanism by which *Pf*A-M17 functions under different concentrations of both protein and metal cofactor. As a result, we have termed the L13 loop (residues 375–401) the ‘regulatory loop’.

**Table 2.**
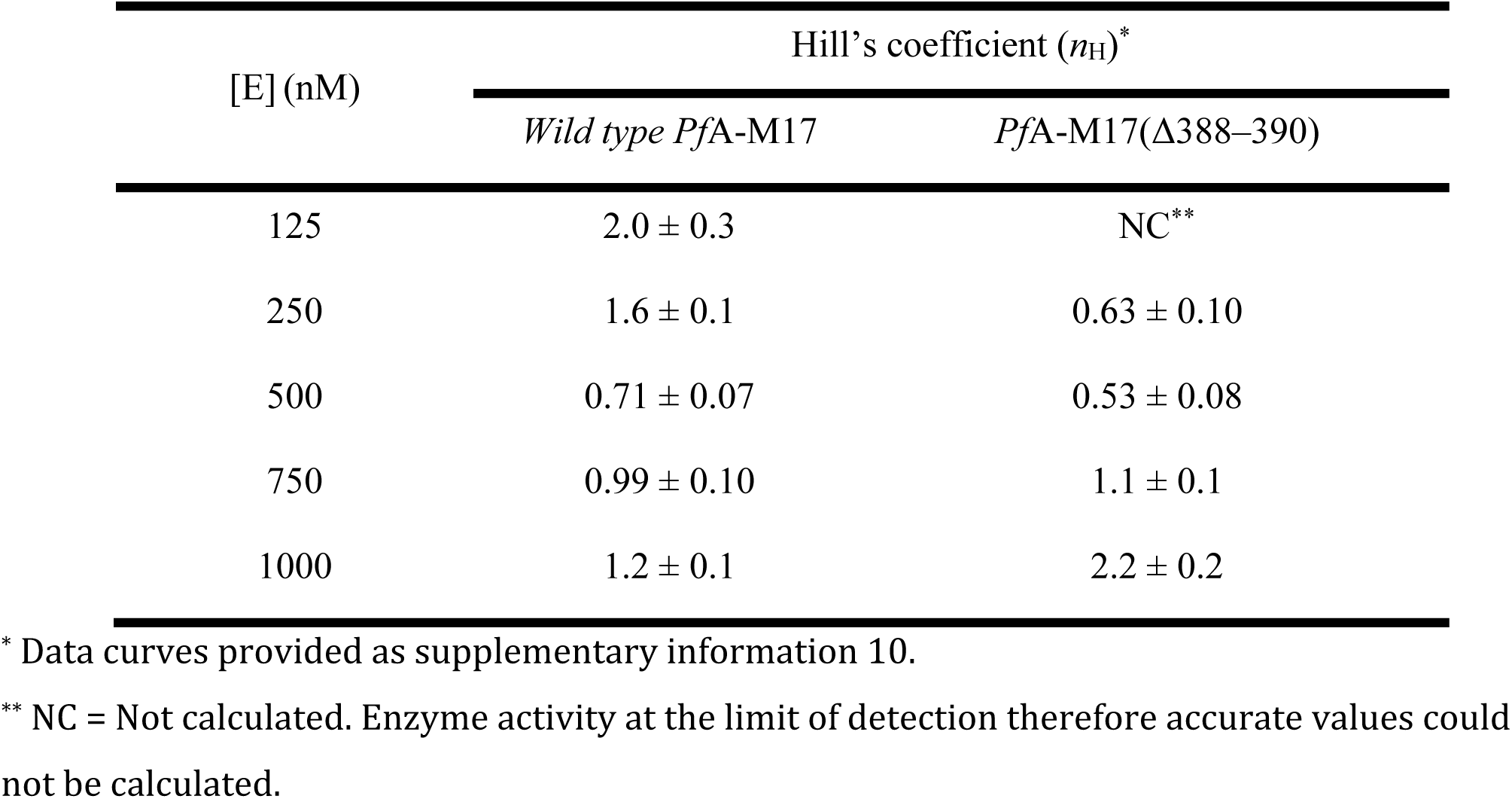
Hill’s coefficient describing the cooperativity of binding of the Mn^2+^ cofactor to *Pf*A-M17 and *Pf*A-M17(Δ388–390).

| [E] (nM) | Hill's coefficient ($n_H$)<sup>*</sup> | |
| --- | --- | --- |
| | *Wild type PfA*-M17 | *PfA*-M17(Δ388–390) |
| 125 | $2.0 \pm 0.3$ | NC<sup>**</sup> |
| 250 | $1.6 \pm 0.1$ | $0.63 \pm 0.10$ |
| 500 | $0.71 \pm 0.07$ | $0.53 \pm 0.08$ |
| 750 | $0.99 \pm 0.10$ | $1.1 \pm 0.1$ |
| 1000 | $1.2 \pm 0.1$ | $2.2 \pm 0.2$ |

<sup>\*</sup> Data curves provided as supplementary information 10.
<sup>\*\*</sup> NC = Not calculated. Enzyme activity at the limit of detection therefore accurate values could not be calculated.

### Regulatory loop differs throughout M17 aminopeptidase family

M17 aminopeptidases play vital roles in a wide range of physiological processes throughout all kingdoms of life. We therefore sought to determine whether the M17 enzymes from other organisms might operate via similar mechanisms to what we have determined for *Pf*A-M17. We performed a sequence alignment of M17 aminopeptidases from a range of different organisms and compared the region of the regulatory loop (Supp. Info. 11). The alignment showed that while Lys386 is highly conserved throughout the M17 aminopeptidase family, neither the regulatory loop following it, nor the aspartic acid residue that links the regulatory loop with the third metal binding site are conserved. Closer examination of the alignment showed that the equivalent loops in M17 aminopeptidases can be classified into two groups. Those like *Pf*A-M17 that possess loops of similar length and nature as well as the linked site 3 (Asp or Glu), and those that have a shortened loop and absence of site 3 (though the site is still conserved in some of these enzymes). Based on the results of our own mutational analyses, we expect enzymes in this latter group are incapable of the loop dynamics described for *Pf*A-M17. The sequence analysis therefore suggests that two distinct mechanisms of proteolytic regulation exist within the family of M17 aminopeptidases.

## Discussion

Our appreciation of protein flexibility and dynamics has grown remarkably from the early static lock-and-key model, to our current understanding that proteins are dynamic entities capable of extreme flexibility. For enzymes, flexibility influences substrate binding (50, 51) and reaction mechanism (52, 53), allowing precise control of reactions that if left unregulated, could be harmful to the cell. Despite the clear importance of enzyme flexibility, there are few cases where dynamics, and how those dynamics link to function, are truly understood on an atomic level.

M17 aminopeptidases were first characterized as homohexamers during early examinations of the blLAP (54). The conserved arrangement is now one of the most distinctive features of the family, along with the arrangement of the binuclear metal center (4). It is therefore somewhat surprising that almost 50 years on we do not understand the contribution the quaternary structure makes to enzyme function. Herein, we hypothesized that the hexameric assembly exerts its effect on function through protein dynamics. Indeed, our analyses of the MD simulations of inactive, monomeric enzyme compared to active, hexameric enzyme, have shown this to be the case. We determined that hexamerization enables catalysis by stabilizing the protomers, thus shielding the essential nucleophilic water. Such a mechanism of preserving active site stability by oligomerization has been previously described, for example, dihydrodipicolinate synthase (DHDPS), which utilises a tetrameric arrangement to stabilize the catalytic dimerization interface for optimal catalytic efficiency (55). Further, formation of the DHDPS tetramer is promoted by substrate binding (56), thus, similarly to *Pf*A-M17, DHDPS demonstrates a complex interplay between substrate-binding, oligomerization, and catalysis that is mediated by protein dynamics.

Beyond a role in active site stability, the hexameric arrangement of *Pf*A-M17 also allows communication between neighbouring active sites through the movement of a dynamic regulatory loop. While first identified in the MD simulations of the hexameric enzyme, we were able to validate the motion using X-ray crystallography, whereby we captured an inactive conformation of the enzyme. In this conformation, the three active sites of each trimer are linked by regulatory loops (L13 loops, one contributed from each protomer), and the binuclear metal center has undergone structural rearrangement. Our mutational analysis of the regulatory loop and metal site 3 demonstrated that loop movement is intrinsically coupled with rearrangement of the binuclear metal center, and that these metallo-protein dynamics are an inherent component of *Pf*A-M17 catalytic function. This does not appear to be a result of changes to the hexameric assembly, but rather the ability of the regulatory loop to mediate communication between the active sites. Therefore, based on the biochemical and structural results, we propose a model in which the transition between the active and inactive conformations of *Pf*A-M17 is part of a dynamic regulatory mechanism that has evolved to moderate the rate of aminopeptidase activity under different environmental conditions.

To fully characterise how the dynamic regulatory loop moderates catalysis, the role of the highly conserved active site lysine residue (Lys386), which is shuttled between the active sites of neighbouring chains, must be considered. This residue has previously been proposed to stabilize the substrate in the active site, but also to act as the catalytic base in the reaction mechanism (16, 44, 46). However, here we demonstrated that the function of Lys386 extends beyond catalysis, and includes a role in hexamer stabilization. Our MD simulations showed that hexamerization influences active site stability. Therefore, it is feasible that the inverse is also true, and that loss of Lys386 affects hexamerization via destabilization of the active site. Genna *et al*. recently reported a comprehensive study comparing the structures and active-site dynamics of a diverse range of DNA and RNA processing enzymes that function via a binuclear metal center (57). The analysis resulted in the identification of a structurally conserved active site architecture wherein basic amino acids occupy conserved positions in relation to the binuclear metal center, and interact with the metals through the second coordination shell (57). Further, the structurally conserved residues all interact with substrate, despite hailing from a diverse range of enzyme families. Importantly, other enzymes with functions beyond DNA processing have described similar architecture, including the hexameric oxylate decarboxylase (58). The parallels between the architecture described by Genna *et al*. (57) and those for the M17 aminopeptidases are striking. The distance between Lys386 of *Pf*A-M17 and the metal center in both the active and inactive conformations (∼ 4 Å) is consistent with occupation of the second shell of metal ions, and Lys386 likely interacts with substrate. How then, does mutation of Lys386 cause dissociation of the hexamer? In the tomato LAP-A1, mutation of the metal-coordination residues as well as the conserved lysine results in hexamer destabilization (59). Therefore, we propose that Lys386 stabilizes the binuclear center through interactions with the second coordination shell, and that in its absence, in *Pf*A-M17(K386A), the binuclear center is destabilized, which in turn disrupts hexamerization.

Further parallels can also be made between the *Pf*A-M17 system and the diverse enzymes that operate via the conserved two-metal ion mechanism (with associated basic residue). A particularly striking similarity is observed between the human DNA Polymerase-η (Pol-η) and *Pf*A-M17; Pol-η operates via a highly dynamic and cooperative process that involves transient recruitment of a third metal ion (60). The conserved basic residue of Pol-η, Arg61, is highly flexible, and specific conformations of Arg61 (sampled as part of a large equilibrium ensemble) serve to recruit incoming substrate, facilitate the reaction, and act as an exit shuttle (60). Such a role could readily be envisaged for Lys386. This is particularly feasible given the flexibility of the loop region (suitable for guiding substrate entry and product exit) and the proposed role in substrate positioning (for reaction facilitation). How then, might this align with the observation that other M17 aminopeptidases do not possess the regulatory loop? While the shortened loops of other M17 aminopeptidases would not be expected to link neighbouring active sites, the conserved lysine would remain capable of stabilizing the binuclear center and facilitating the catalytic reaction, and additionally, the shortened loop would still be expected to be capable of the flexibility required to chaperone substrate entry and/or product exit. Therefore, the mechanism of the reaction, and role of Lys386 in facilitating that reaction, would be expected to remain conserved between M17 aminopeptidases, but not the cooperative metal-dependent switch by which the *Pf*A-M17 reaction is regulated.

There is already some suggestion in the literature that the regulatory mechanisms of M17 aminopeptidases differ. For example, the M17 aminopeptidase from *Helicobacter pylori*, which does not possess the flexible regulatory loop, has been shown to exhibit positive cooperativity with the binding of a fluorescent reporter substrate (61). To our knowledge, this cooperativity has not been observed for any other M17 aminopeptidase. Given that the diverse family of M17 aminopeptidases operate within a wide range of different organisms, it is entirely reasonable to expect that the enzymes have evolved to operate differently within diverse environments. Indeed, it has been suggested that flexible loops in proteins can facilitate the emergence of novel functions, which results in divergent evolution (62-64). In terms of *Plasmodium* parasites, the concentrations of different divalent metal cations are known to fluctuate throughout the life cycle (65). Therefore, the metal-dependent switch described here could potentially serve to moderate the rate of proteolysis to suit different metabolic demands throughout the complex parasite life cycle(13, 14).

The implication of the two-metal center re-arrangement, which includes a third metal-binding site, should also be considered further. The metal sites of M17 aminopeptidases are most often occupied by zinc ions (Zn^2+^); however, can also be exchanged for other metals such as cobalt (Co^2+^), manganese (Mn^2+^), magnesium (Mg^2+^), and copper (Cu^2+^) (4). Previous examinations have focused on assigning identity and role to each of the site 1 and site 2 metal ions, with each of them considered to have distinct catalytic roles (regulatory vs catalytic). It has never before been considered that the metal positions themselves might change. The finding that the binuclear site is dynamic, and that the site 3 metal ion plays a role in the catalytic cycle of *Pf*A-M17, therefore carries substantial implications for previous and future work. In crystal structures, M17 aminopeptidases have often been observed with only a single metal ion bound. These one-metal ion forms of the enzyme have been used to interpret kinetic examinations; biphasic curves that result from titration of metal chelators into *Pf*A-M17 have been interpreted as two-metal ion bound (100% activity), one-metal ion bound (∼ 50 % activity), and unbound (irreversibly inactivated) forms of the enzyme (47). However, our characterization of a third metal binding site, which is sampled as part of the catalytic reaction, casts doubt on that interpretation. In a separate line of study, we have recently characterized a novel mode of *Pf*A-M17 inhibition, whereby small molecule inhibitors reversibly displace the site 2 ‘catalytic’ metal (24). Furthermore, studies of a λ-exonuclease, which possesses the conserved architecture discussed above (two-metal ion mechanism with associated basic residue), proposed transient binding of a third metal ion, and additionally, demonstrated that the timing of both coordination and dissociation of each of the binuclear metal-ions is important to the catalytic cycle (66). Therefore, *Pf*A-M17 could potentially transition between a mono-, bi-, and tri-metal ion center as part of the catalytic cycle.

Taking all of this into account, we propose a novel mechanism for the regulation of *Pf*A-M17 activity through interplay of metallo-protein regulatory dynamics. We suggest that *Pf*A-M17 exists in a dynamic equilibrium between an inactive conformation, which does not possess the active site arrangement to accept substrate nor perform catalysis, and an active conformation, capable of product turnover. Transition between these conformations is mediated by the cooperative movement of six regulatory loops (one contributed from each unit of the hexamer), which, through re-positioning Lys386, mediate rearrangement of the binuclear metal center. Finally, we propose this transition occurs in response to specific environmental signals, which act to regulate the rate of *Pf*A-M17 catalysis according to physiological demands of the *Plasmodium* parasite.

## Conclusion

The M17 aminopeptidases have been of interest as key players in a wide range of physiological processes for over 50 years. From a protein mechanics perspective, the conserved hexameric assembly, containing six discrete active sites with binuclear metal centers, represents an exciting system in which to investigate the role of dynamic cooperativity and function within a large oligomer. Herein, we have shown that the hexameric assembly is absolutely essential for the proteolytic activity of *Pf*A-M17. The arrangement stabilizes and preserves key catalytic machinery within the six active sites, and further, plays a role in the regulation of aminopeptidase activity. Within the hexamer, each of the two disc-like trimers operate independently; a flexible loop links each of the three active sites within each trimer to its neighbour, thereby operating cooperatively to convert the enzyme from the inactive to active state. Further, movement of the regulatory loop is coupled with a rearrangement of the binuclear metal center. Our studies show that not only do metal ion dynamics exist within the active site, but that the positions of the metals are manipulated to control the activity of the enzyme. We therefore propose the dynamic transition between inactive and active states is part of a key regulatory mechanism that controls the activity of *Pf*A-M17, and that the transition is moderated by changes in the physiological environment.

## Abbreviations

*Pf*A-M17: *Plasmodium falciparum* M17 aminopeptidase
AUC: analytical ultracentrifugation
MD: molecular dynamics
PCA: principle component analysis
RDF: radial distribution function

## Author Contributions

S.M. and N.D. designed the research; N.D., W.Y., K.K.S., B.T.R., S.C.A., T.R.M, B.K.H, and I.K. performed the research and analysed the data; A.M.B. and S.M. supervised the research; N.D. and S.M. wrote the manuscript with contributions from all other authors.

## Acknowledgements

We thank the National Health and Medical Research Council (Project Grant 1063786 to S.M. and P.J.S.; Fellowship 1022688 to A.M.B.) and the Australian Research Council (Fellowship FT100100690 to S.M.) for funding support. This work was supported by the Victorian Life Sciences Computation Initiative (VLSCI) and National Computational Infrastructure (NCI). This research was undertaken in part using the MX2 beamline at the Australian Synchrotron, part of ANSTO, and made use of the ACRF detector.

## Associated content

The following figures and movies are provided as supplementary information:

**Supplementary Information 1.** Circular dichroism analysis of *Pf*A-M17.

**Supplementary Information 2:** Analytical ultracentrifugation of *Pf*A-M17.

Supplementary Information 3: RMSD plot of monomeric and hexameric *Pf*A-M17 over course of molecular dynamics simulations.

**Supplementary Information 4:** Radial distribution function (RDF) plot of water molecules in the zinc environment from the MD simulations of monomeric and hexameric *Pf*A-M17.

**Supplementary Information 5:** RMSF analysis throughout molecular dynamics simulations.

**Supplementary Information 6:** Movie showing displacement along PC2 of *Pf*A-M17 hexamer simulation.

**Supplementary Information 7:** Crystallographic Data Collection and Refinement Statistics.

**Supplementary Information 8:** Movie showing morph between active and inactive *Pf*A-M17.

**Supplementary Information 9:** Analytical size exclusion chromatography of *wild type* and mutant *Pf*A-M17.

**Supplementary Information 10:** Cooperativity on metal binding of *wild type Pf*A-M17 compared to *Pf*A-M17(Δ388-390).

**Supplementary Information 11:** Sequence alignment of M17 aminopeptidases from a range of organisms.

## Table of Contents Graphic

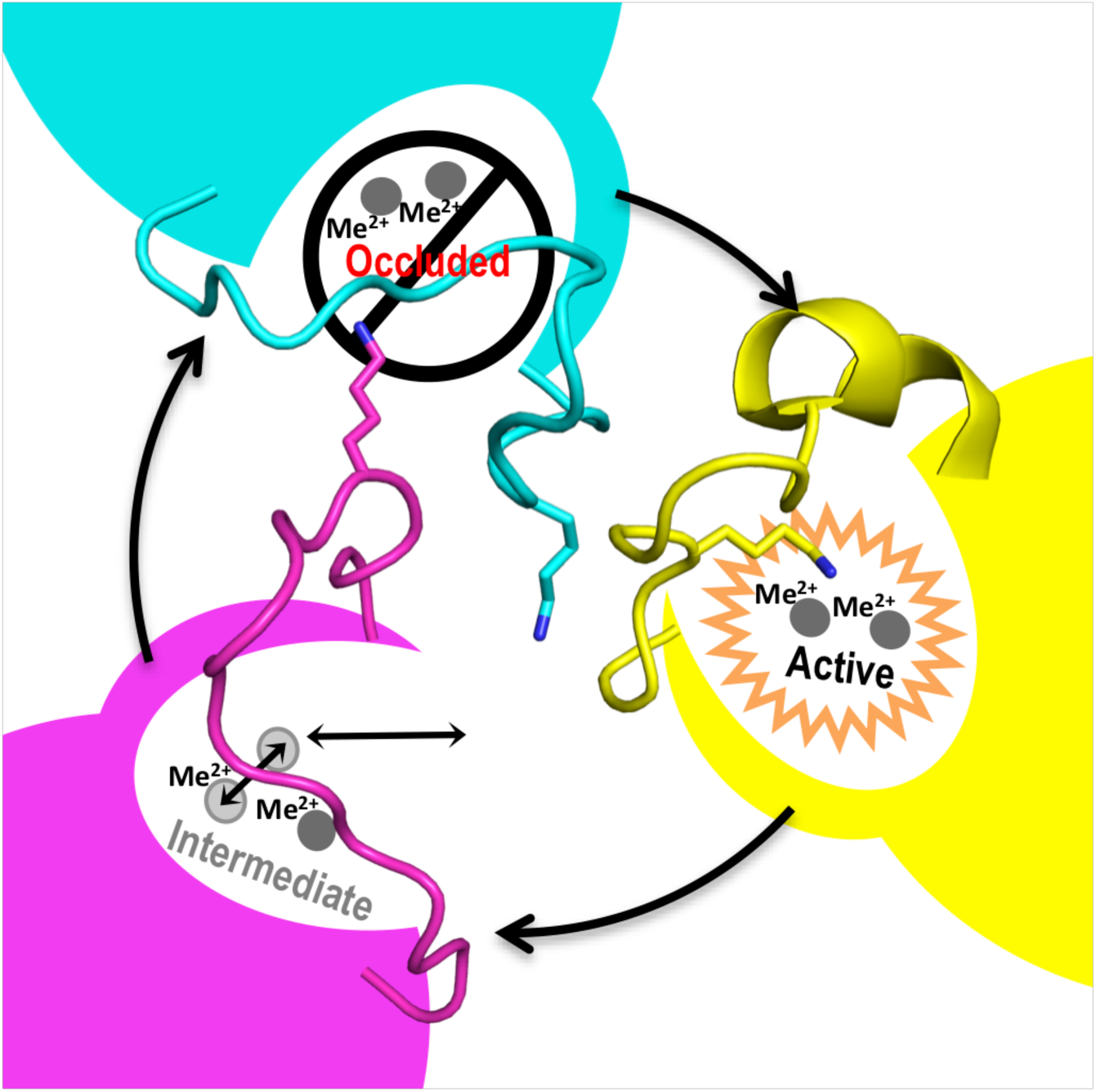

