## Supplementary Material for "A metal-dependent switch moderates activity of the hexameric M17 aminopeptidases"

**Supplementary Information 1: Circular dichroism analysis of *Pf*A-M17.** Comparison of circular dichroism profiles of *wild type Pf*A-M17 (blue) and *Pf*A-M17(W525A+Y533A) (red) show that they have comparable secondary structure composition.

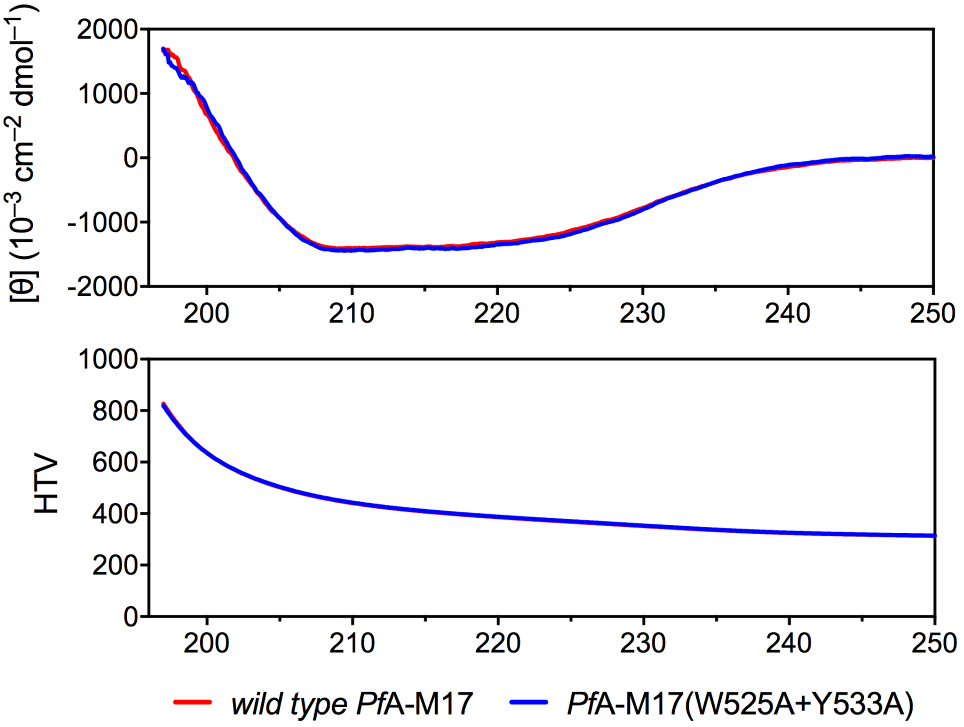

**Supplementary Information 2: Analytical ultracentrifugation of *wild type Pf*A-M17 and *Pf*A-M17(W525A+Y533A).** Analysis of sedimentation velocity data of *wild type Pf*A-M17 (black) and *Pf*A-M17(W525A+Y533A) (grey) in 1.0 mM $MnCl_2$ represented as (A) continuous $c(s)$ distribution and corresponding residuals and (B) continuous c(M) distribution and corresponding residuals. Global fit analysis of *Pf*A-M17(W525A+Y533A) at 8.6 µM (circle), 4.3 µM (triangle) and 1.7 µM (diamond) and corresponding residuals at two different speeds (C) 12,000 rpm and (D) 18,000 rpm indicate that *Pf*A-M17(W525A+Y533A) is a monomer in solution. (E) Hydrodynamic properties of *Pf*A-M17.

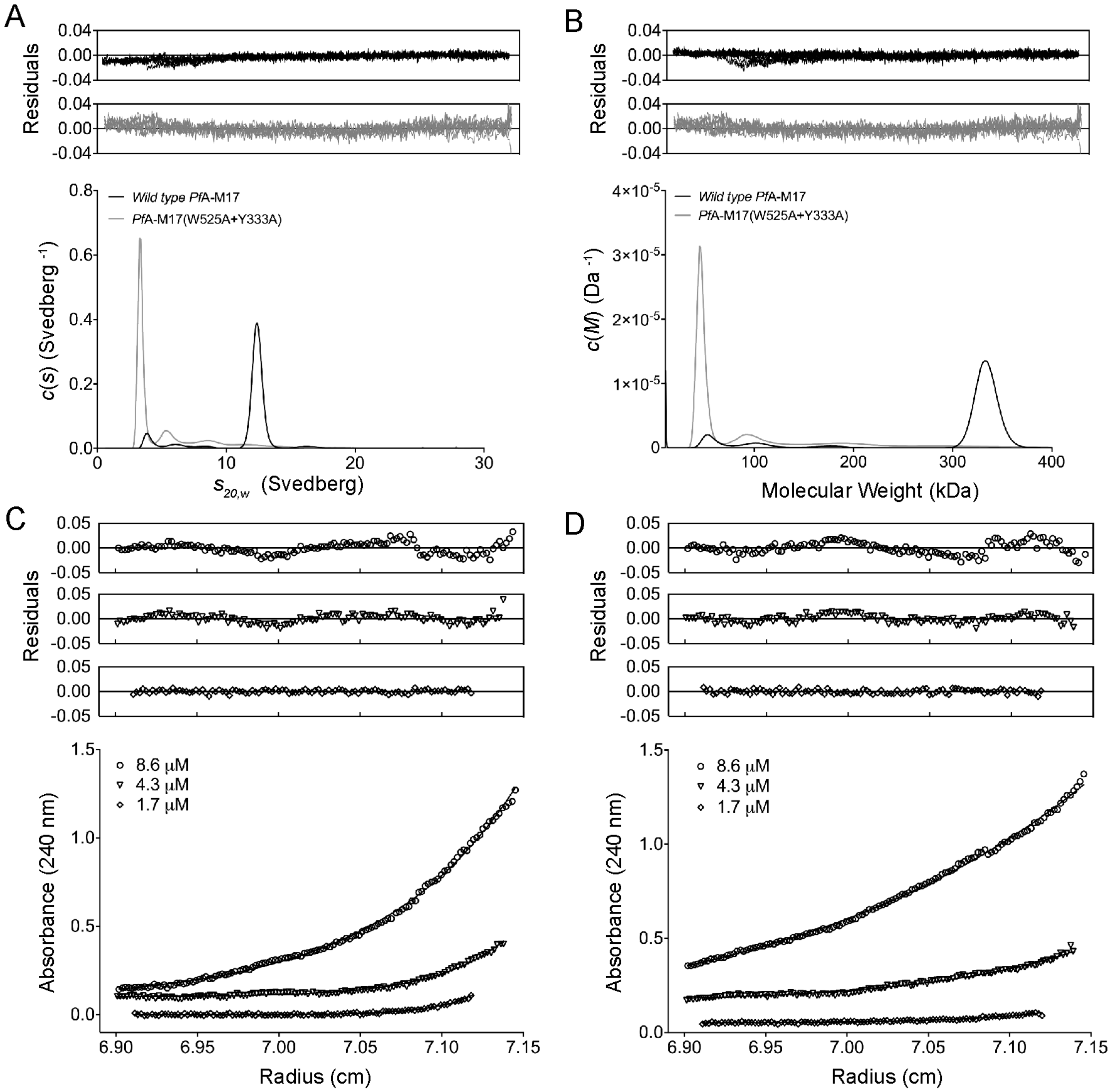


E

**Hydrodynamic properties of *Pf*A-M17 calculated by AUC**

| | $M_r$ [a] | $s_{20,w}$ [b] | $M_1$ [c] | $M_2$ [d] | $f/f_0$ [e] |
|---|---|---|---|---|---|
| ***Pf*A-M17** | 352 | 12.4 | 332 | – | 1.01 |
| ***Pf*A-M17(Y533A+W525A)** | 58.6 | 3.3 | 56.0 | 59.4 | 1.45 |

[a] Relative molecular weight calculated from the amino acid sequence (kDa).

[b] Standardized sedimentation coefficient taken from the ordinate maximum of the $c(s)$ distribution best fits (panel A).

[c] Molar mass determined from the ordinate maximum of $c(M)$ distribution best fits (Panel B).

[d] Molar mass determined from sedimentation equilibrium data fit to a single species model (panels C and D)

[e] Frictional coefficient calculated from $s_{20,w}$ using the $\bar{v}$ method employing SEDNTERP

**Supplementary Information 3:** RMSD plot of (A) monomeric, and (B) hexameric, *Pf*A-M17 over course of triplicate molecular dynamics simulations.

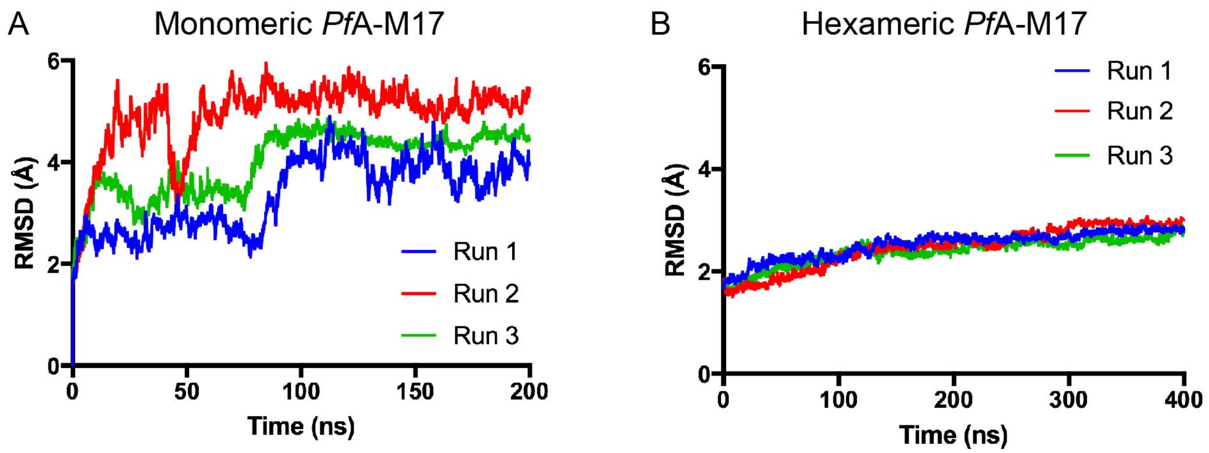

**Supplementary Information 4: Radial distribution function (RDF) plot of water molecules in the zinc environment from the MD simulations of (A) monomeric *Pf*A-M17, and (B) hexameric *Pf*A-M17.** The RDF was calculated by the average of all six sites in the hexamer. Probability percentage was calculated by the occupancy of the water divided by the number of waters (as measured by oxygen atom of the waters) at any given distance to the zinc ions. The plots show that the catalytic water in simulation of monomer is unstable, however, the hexamer has a single water molecule in the catalytic position throughout the simulation.

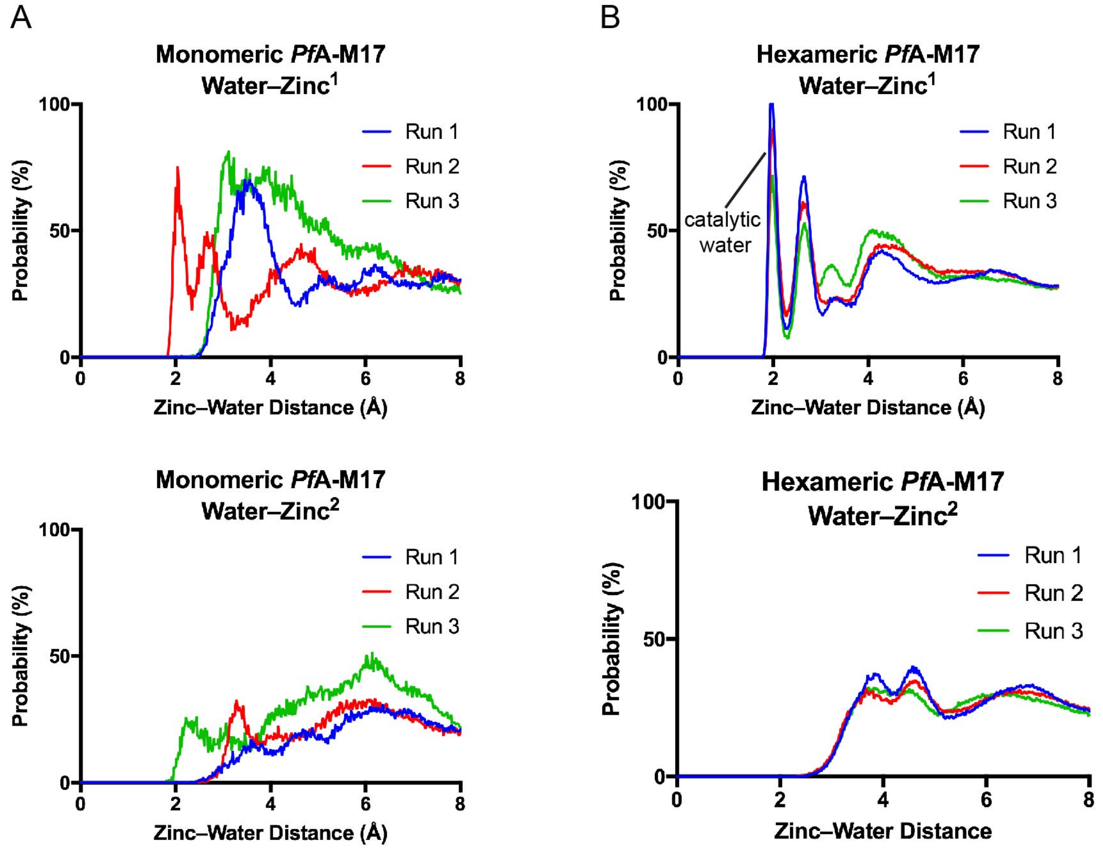

**Supplementary Information 5: RMSF analysis throughout molecular dynamics simulations.** (A) *Pf*A-M17 monomer throughout simulation. (B) RMSF analysis of the six individual of hexameric *Pf*A-M17 throughout simulation.

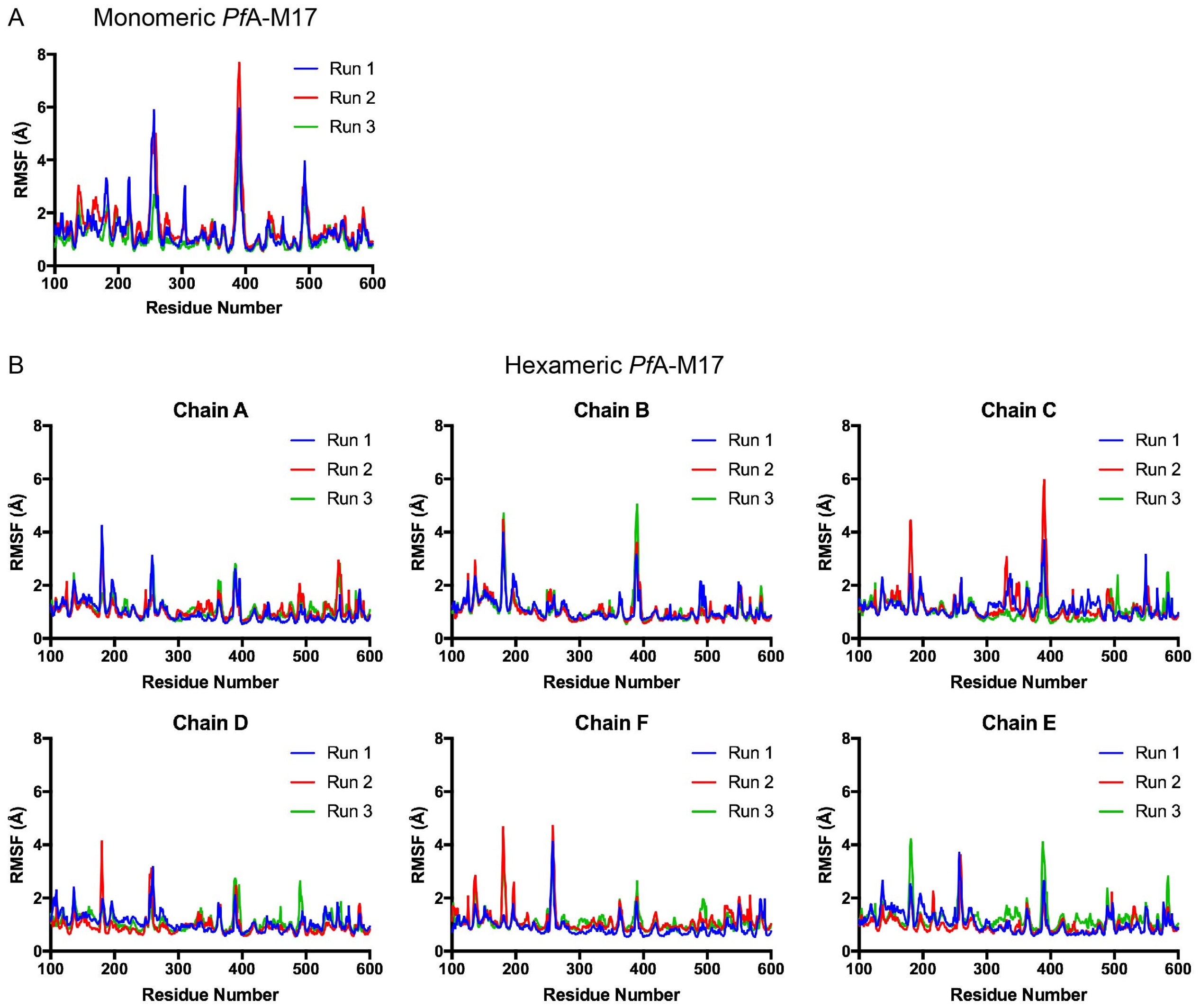

**Supplementary Information 6: Movie showing displacement along PC2 of *Pf*A-M17 hexamer simulation.** *Pf*A-M17 trimer with greatest level of movement shown only (chains D, E, and F), with chain D in yellow, E in red, and F in orange.

https://www.dropbox.com/s/2mjkvwokzqnz2df/SupMov7_2880x2160.10Mbps.mp4?dl=0

### Supplementary Information 7: Crystallographic Data Collection and Refinement Statistics

| | |
|---|---|
| **Data collection** | |
| Space Group | P 2 21 21 |
| Cell dimensions: | |
| *a, b, c* (Å) | 111.54, 172.67, 179.67 |
| α = β = γ (°) | 90.0 |
| Resolution range (Å) | 49.19 – 2.00 (2.03 – 2.00) |
| Total Observations | 1,626,188 (82,021) |
| Unique Observations | 233,061 (11,469) |
| Multiplicity | 7.0 (7.2) |
| Completeness (%) | 100.0 (100.0) |
| <I/σI> | 7.0 (0.7) |
| CC(1/2) | 0.997 (0.244) |
| $R_{pim}$ (%) | 7.1 (125.5) |
| PDB | 6NWI |
| **Structure refinement** | |
| *Non hydrogen atoms* | |
| Protein | 23594 |
| Water | 1669 |
| Ligand | 128 |
| $R_{work}$ *(%)* | 19.1 |
| $R_{free}$ *(%)* | 22.5 |
| *RMS deviations* | |
| Bond lengths (Å) | 0.004 |
| Angles (°) | 0.50 |
| *Ramachandran plot* | |
| Favoured(%) | 97.0 |
| Outliers (%) | 0.03 |
| *B factors* ($Å^2$) | |
| Protein | 39.2 |
| Ligands | 45.5 |
| Solvent | 42.9 |

**Supplementary Information 8: Movie showing morph between active and inactive *Pf*A-M17.**
Chain A in blue, B in purple, C in teal, D in yellow, E in red, and F in orange.

https://www.dropbox.com/s/h6nvknk7asrfjcz/SupMov9_2880x2160.10Mbps.mp4?dl=0

**Supplementary Information 9: Analytical size exclusion chromatography of *wild type* and mutant *Pf*A-M17.** Absorbance traces of analytical gel filtration experiments to estimate oligomeric state of variant *Pf*A-M17 enzymes. Dashed lines indicate predicted elution volumes of different oligomeric states. Oligomeric state of *Pf*A-M17 (red, hexameric) and *Pf*A-M17(W525A+Y533A) (magenta, monomeric), confirmed by AUC. Based on interpolation of standard curve, elution volume of hexameric *Pf*A-M17 (red) corresponds to an approximate molecular weight of 330 kDa (molecular weight of hexameric *Pf*A-M17 calculated from amino acid sequence is 352 kDa), whereas elution volume of monomeric *Pf*A-M17(W525A+Y533A) (magenta) corresponds to approximate molecular weight of 68 kDa (monomeric *Pf*A-M17 molecular weight from amino acid sequence is 58.6 kDa). *Pf*A-M17(K386A) (orange) elution volume corresponds to an approximate molecular weight of 143 kDa, while *Pf*A-M17(D394A) (light blue), *Pf*A-M17(A387P) (dark blue), *Pf*A-M17(Δ388-389) (grey), and *Pf*A-M17(Δ388-390) (green) are all largely hexameric.

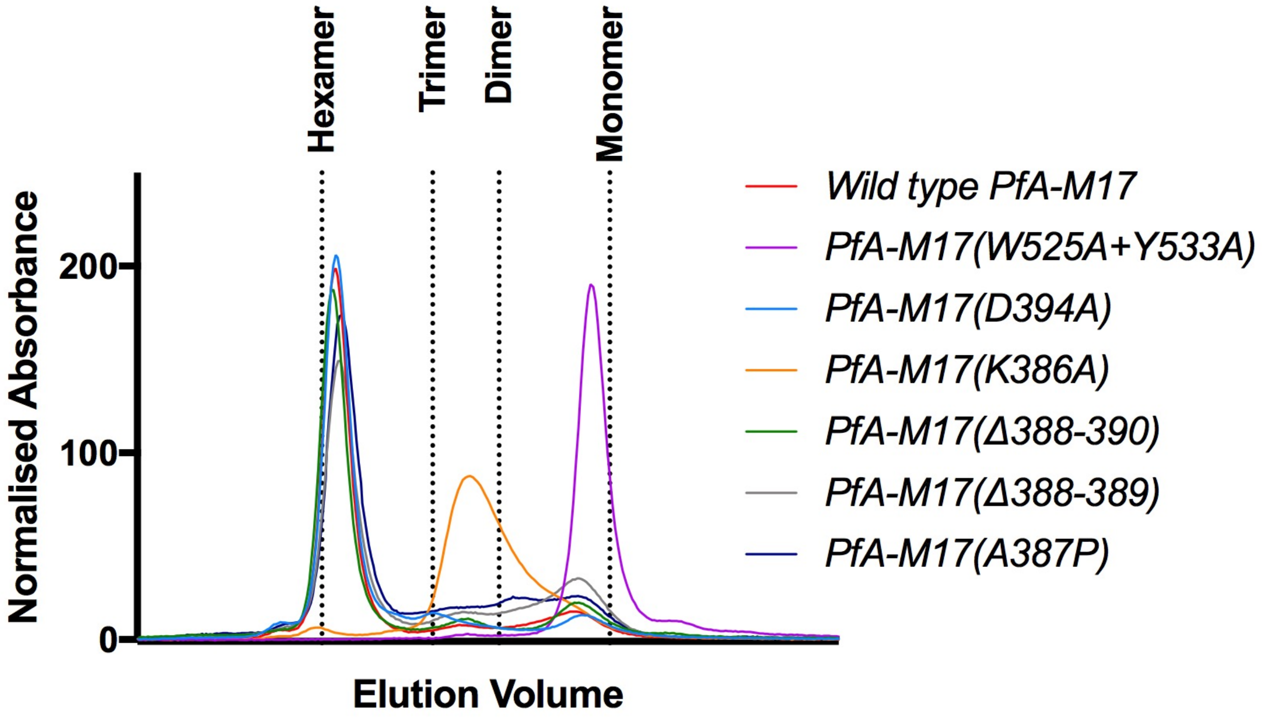

**Supplementary Information 10: Cooperativity on metal binding of *wild type Pf*A-M17 compared to *Pf*A-M17(Δ388-390).** (A) titration of $Mn^{2+}$ (10 μM – 5/10 mM) into wild type *Pf*A-M17 (125 – 1000 nM). Data normalised for percentage activity.

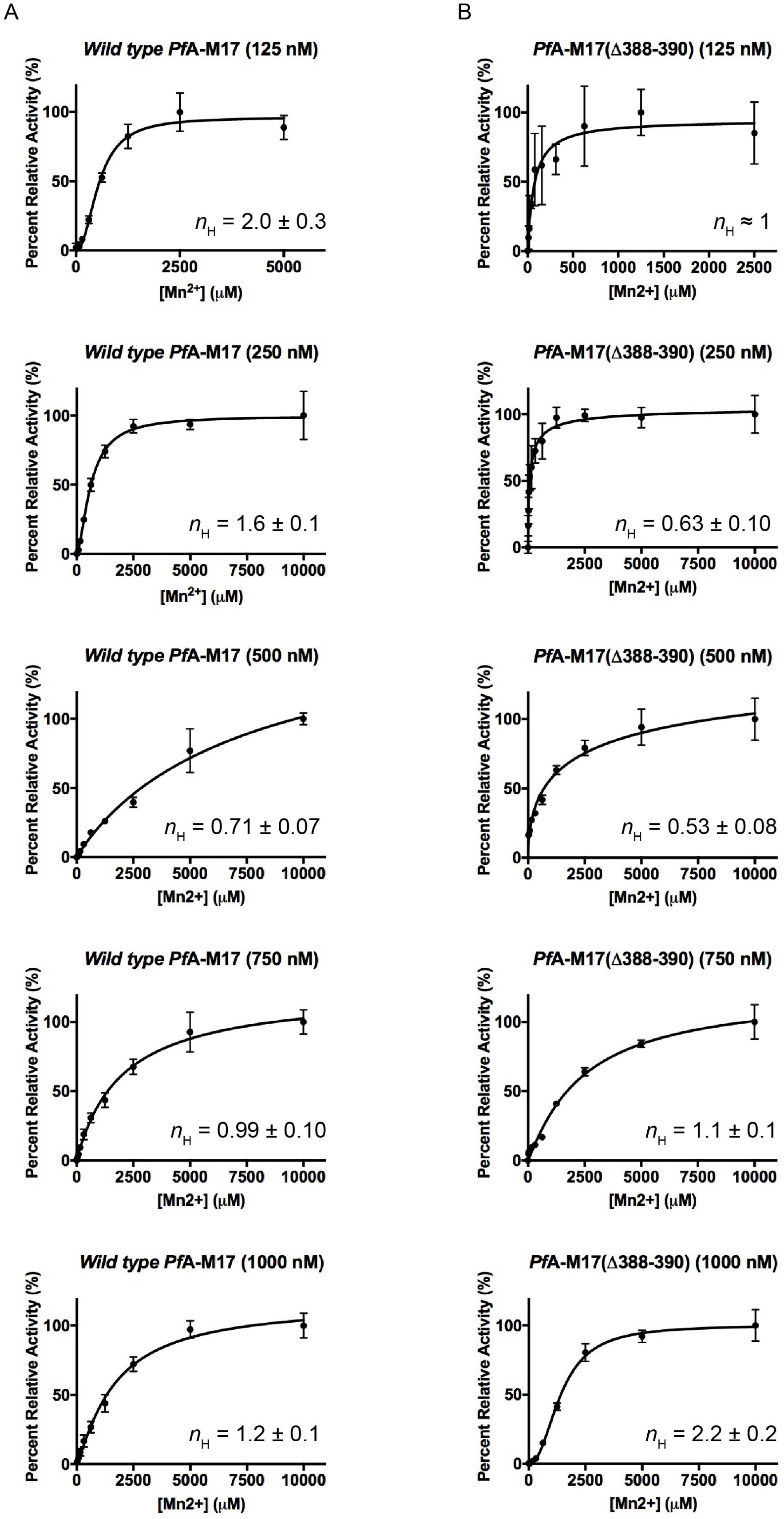

**Supplementary Information 11: Sequence alignment of verified M17 aminopeptidases from a range of organisms**. Conserved lysine residue (Lys386 in *Pf*A-M17) is highlighted in green and loop residues are highlighted in pink. Yellow highlight shows that Asp394 of *Pf*A-M17, representative of zinc binding site 3, is not conserved throughout the enzyme family. Blue box indicates the M17 aminopeptidases that possess a loop of similar length and nature to *Pf*A-M17. Alignment performed with Clustal Omega.

```
C. elegans                YHVGK-AG--PTPPAFVVLSHEVP-G-STEHIALVGKGVVYDTGGLQIKT--KTGMPNM   272
B. taurus                 LSVAK-GS--EEPPVFLEIHYKGSPNASEPPLVFVGKGITFDSGGISIKA--AANMDLM   296
H. sapiens                LSVAK-GS--DEPPVFLEIHYKGSPNANEPPLVFVGKGITFDSGGISIKA--SANMDLM   302
M. musculus               LSVAK-GS--EEPPVFLEIHYMGSPNATEAPLVFVGKGITFDSGGISIKA--SANMDLM   302
R. norvegicus             LSVAK-GS--EEPPVFLEIHYTGSPNATEAPLVFVGKGITFDSGGISIKA--SANMDLM   302
A. gambiae                LTVAK-GS--CEPPIFLELSYYGT-NTKERPIVLIGQGNTFDSGGLCLKT--LEALTDM   315
A. thaliana               LAVAA-AS--ANPPHFIHLIYKPSSGPVKTKLALVGKGLTFDSGGYNIKTGPGCLIELM   373
S. lycopersicum LAP-N     LRVAA-AS--ANPAHFIHLCYKPSSGEIKKKIALVGKGLTFDSGGYNIKTGPGCSIELM   361
S. tuberosum              LGVAAAAT--ENPPYFIHLCFKTNSRERKTKIALVGKGLTFDSGGYNLKTGAGSKIELM   345
S. lycopersicum LAP-A     LAVAAAAT--ENPPYFIHLCFKTPTKERKTKLALVGKGLTFDSGGYNLKVGAGSRIELM   364
T. annulata               LSVCQ-GS--KFPPKFLHAVYKP-DGEVKKTLAFVGKGITFDAGGYNVKN-SSTMIHYM   318
T. gondii                 LAVAK-GS--LFPPQFIHLTYKS-SN-PKRRVAFVGKGICFDAGGYNLKT-GGAQIELM   566
P. falciparum             LSVGK-GS--MYPNKFIHLTYKS-KGDVKKKIALVGKGITFDSGGYNLKAAPGSMIDLM   396
C. hominis                LAVAQ-GS--KSPAQFVHLTYKP-KGEIKKRIALVGKGITMDTGGYNIKH---QMIHFM   315
C. parvum                 LAVAQ-GS--KSPAQFVHLTYKP-KGEIKKRIALVGKGITMDTGGYNIKH---QMIHFM   341
L. major                  YNVGR-GS--RYEPYLVVLEYIGNPR-SSATTAIVGKGVTFDCGGLNIKP--YTSMETM   345
H. pylori                 LAVNK-ASLSVNPPRLIHLVYKPKKA--KKKIALVGKGLTYDCGGLSLKP--ADYMVTM   284
S. aureus                 HAVGK-GS--EHPPVVITMTYNGGDS-NDAPIALVGKGITYDSGGYSIKS--KIGMQTM   280
E. coli                   LAVGQ-GS--QNESLMSVIEYKGNASEDARPIVLVGKGLTFDSGGISIKP--SEGMDEM   290
R. typhi                  LGVGQ-GS--QNESKLVVMEYKGGSR-DDSTIALVGKGVIFDTGGISLKP--SSNMHLM   285
C. tetani                 LAVSK-GS--FEESQLIVMNYKGNSN-SDKKLALVGKGLTYDSGGYSIKP--TSSMINM   267
B. cereus                 LAVNQ-GS--VEPPKMIALIYKGKEE-WKDVIGFVGKGITYDTGGYSLKP--REGMVGM   280
                            *   .        .    .            ::*:*   * **  :*      :  *
```
